# Dual clathrin and adhesion signaling systems regulate growth factor receptor activation

**DOI:** 10.1101/2020.11.09.373837

**Authors:** Marco A. Alfonzo-Mendez, Kem A. Sochacki, Marie-Paule Strub, Justin W. Taraska

## Abstract

The crosstalk between growth factor and adhesion receptors is key for cell growth and migration. In pathological settings, these receptors are drivers of cancer. Yet, how growth and adhesion signals are spatially organized and integrated is poorly understood. Here we use quantitative fluorescence and electron microscopy to reveal a mechanism where flat clathrin lattices partition and activate growth factor signals via a coordinated response that involves crosstalk between epidermal growth factor receptor (EGFR) and the adhesion receptor β5-integrin. We show that ligand-activated EGFR, Grb2, Src, and β5-integrin are captured by clathrin coated-structures at the plasma membrane. Clathrin structures dramatically grow in response to ligand activation into large flat plaques and provide a signaling platform that link EGFR and β5-integrin through Src-mediated phosphorylation. Disrupting this EGFR/Src/β5-integrin axis prevents both clathrin plaque growth and receptor signaling. Our study reveals a reciprocal regulation of clathrin lattices and two different receptor systems to enhance cell growth factor signaling. These findings have broad implications for the control of growth factor receptors, mechanotransduction, and endocytosis.

## INTRODUCTION

Cellular communication begins with a cascade of molecular interactions initiated by plasma membrane (PM) receptors of which the receptor tyrosine kinases (RTKs) are one of the major superfamilies. RTKs are ubiquitous integral membrane proteins in eukaryotes that perform numerous actions including the regulation of cell proliferation, differentiation, survival and migration^1^. Epidermal growth factor receptor (EGFR) is one of the most widely studied members of the RTK superfamily, which regulates epithelial tissue development and homeostasis. In lung, breast, and head and neck cancers, EGFR is a driver of tumorigenesis and is a major target for therapy^2, 3^. EGFR spans the membrane and contains a ligand-interacting domain facing the extracellular space and a tyrosine kinase region in the cytoplasm^4^. The binding of EGFR to epidermal growth factor (EGF) triggers receptor dimerization and cross-phosphorylation of tyrosine residues within its cytosolic domain^5, 6^. This provides docking sites for the recruitment of scaffold proteins including Grb2 and the activation of downstream tyrosine kinases such as Src^7–9^. The EGFR pathway can be activated either directly by its cognate ligands or through transactivation by other signaling proteins including integrins^10, 11^.

Integrins are adhesion molecules that are responsible for cell-cell and cell-matrix interactions, and relay mechanical signals bidirectionally between the extracellular space and the cytoplasm^12^. In this way, integrins sense the local environment and control tissue rigidity, cell growth, and movement^13^. Dysregulation of integrin signaling contributes to cancer and metastasis ^14^. Integrins are thought to be heterodimers of α and β subunits, each containing a large multidomain extracellular region (>700 residues) for ligand biding, a single transmembrane helix, and a short cytoplasmic tail (13 - 70 residues)^15^. β-integrin cytoplasmic tails lack enzymatic activity. Instead, they harbor distinct regulatory sequences including two NPxY motifs (Asn-Pro-x-Tyr) that can be phosphorylated^16^. NPxY motifs have a high affinity for phosphotyrosine binding domain proteins^17^. This allows them to bind to signaling partners including clathrin-mediated endocytosis (CME) accessory proteins^18, 19^. While these proteins are known to interact generally, how they are spatially organized at the PM to control signaling and crosstalk is unclear.

EGFR and integrins can be regulated through CME—the main pathway used by eukaryotic cells to internalize receptors into the cytoplasm^20, 21^. During CME, clathrin and adaptors assemble to bend the membrane into Ω-shaped pits^22, 23^. Scission of clathrin-coated pits (CCPs) from the PM yields closed spherical vesicles with an average diameter of ~100 nm^24^. Besides CCPs, cells exhibit a subset of clathrin coats known as flat clathrin lattices (FCLs) or plaques^25, 26^. In contrast to CCPs, FCLs are long-lived on the PM and display a variety of two-dimensional shapes^27, 28^. FCLs are abundant in myocytes and can bind cortical actin during muscle formation and function^29, 30^. FCLs are also important for the adhesion between osteoclasts and bone^31, 32^. During cell communication, different types of receptors including EGFR are clustered at FCLs^33^. Additionally, FCLs are enriched with β5-integrin^34–36^. Yet, it is unknown how flat clathrin lattices are regulated and their general functions are still unclear.

Here, we identified an EGFR/β5-integrin/Src signaling axis that regulates flat clathrin lattice biogenesis during growth factor stimulation. Using a combination of quantitative fluorescence and electron microscopy, we showed that EGF triggers large ultrastructural changes in the membrane of human squamous (HSC3) cells. These changes include the generation and expansion of large FCLs and required EGFR interactions with EGF as well as Src kinase and β5-integrin. Agonist stimulation leads to persistent recruitment of EGFR, Grb2, and β5-integrin into clathrin structures, and a corresponding loss of Src kinase. We provide evidence of β5-integrin phosphorylation mediated by Src that regulates this signaling system. These data reveal a mutual regulation of FCLs and two different receptor systems: EGFR and β5 integrin. Thus, an EGFR/β5-integrin/Src axis contributes to the biogenesis of FCLs, which in turn act as dynamic signaling platforms to partition and enhance growth factor signaling at the PM of human cells.

## RESULTS

### EGF triggers changes in the ultrastructure of clathrin lattices

First, we tracked the ultrastructure of clathrin at the PM during growth factor stimulation. To accomplish this, we used genome-edited HSC3 cells endogenously expressing EGFR tagged with GFP, an established model to study EGFR endocytosis and human EGFR-dependent head and neck carcinoma^37–39^. We treated HSC3 cells with vehicle (Ctrl) or EGF for 2, 5, 15, 30 and 60 min. Then, we mechanically unroofed cells to directly visualized clathrin at the cytoplasmic face of the PM using platinum replica transmission electron microscopy (PREM)^40^. In these images, we measured the structure and distribution of clathrin coated structures (CCSs) across the PM of control and EGF stimulated cells. Remarkably, CCSs were 4.6-fold more abundant in cells stimulated with EGF for 15 min (Fig. 1a-d and Sup. Fig.1).

**Figure 1.**
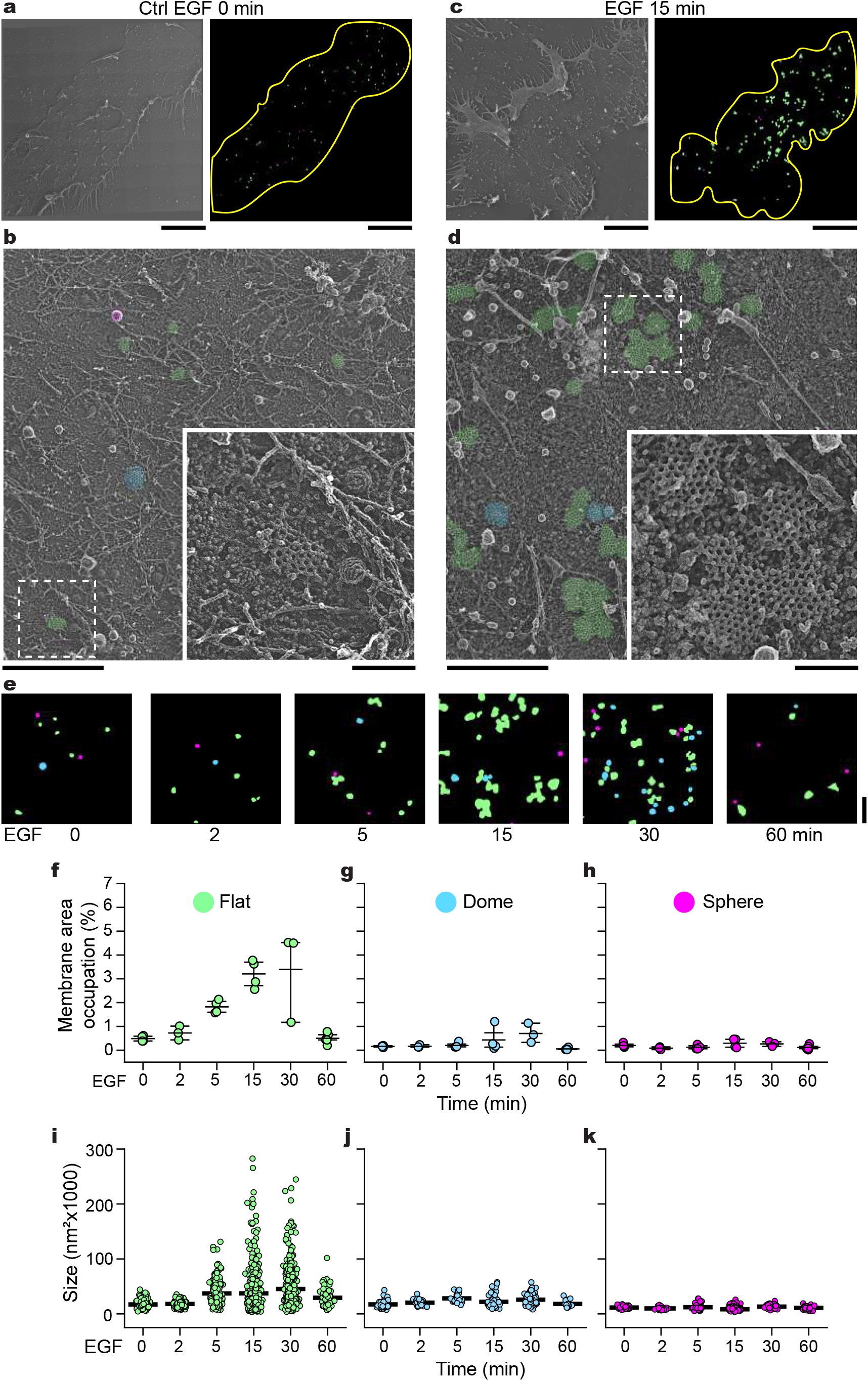
EGF modifies the ultrastructure of clathrin at the plasma membrane. **a**, Montaged PREM image of an unroofed control HSC3-EGFR-GFP cell and the mask created after segmentation of the full membrane outlined (yellow). Flat, dome, and sphere clathrin-coated structures (CCCs) are shown in green, blue, and magenta, respectively. **b**, High-magnification of the cropped PREM in (**a**); the different segmented CCSs are color-coded as in (**a**), with native grayscale in magnified insets. **c**, Montaged PREM image of an unroofed HSC3-EGFR-GFP cell treated with 50 ng/mL EGF for 15 min and the mask created after the segmentation. **d**, High-magnification of a representative region of the PREM in (**c**), the magnification insets are shown at the same scale and are outlined with dashed squares in each image. **e**, Representative masks of the EGF stimulation time course for 0, 2, 5, 15, 30 and 60 min. PREM images corresponding to the masks and cropped images are shown in Supplementary Figure 2. **f-h**, Morphometric analysis of the percentage of plasma membrane (PM) area occupation for each CCS. I-shaped box plots show median extended from 25th to 75th percentiles, and minimum and maximum data point whiskers with a coefficient value of 1.5. **i-k**, Morphometric analysis of the size of flat, dome, and sphere CCSs during the EGF time course. Dot plots show every structure segmented, the bar is the median. 0 min: N_flat_=141, N_dome_=46, N_sphere_=68, N_cells_=4; 2 min: N_flat_=164, N_dome_=32, N_sphere_=30, N_cells_=3; 5 min: N_flat_=184, N_dome_=26, N_sphere_=36, N_cells_=4; 15 min: N_flat_=559, N_dome_=67, N_sphere_=207, N_cells_=4; 30 min:N_flat_=395, N_dome_=149, N_sphere_=113, N_cells_=3; 60 min: N_flat_=81, N_dome_=15, N_sphere_=57, N_cells_=5. Scale bars in (**a**) and (**c**) are 5000 nm. Scale bars in (**b**, **d**, **e**) are 1 μm; insets are 200 nm.

PREM allowed us to segment distinct CCSs based on their curvature into three subclasses: 1) flat clathrin lattices (FCLs), where no curvature is evident (shown in green); 2) dome, curved but the clathrin edge is visible (blue); and 3) sphere, highly curved and edge is not observable (magenta) (Fig. 1a-d)^41^. Representative segmentation masks illustrate the densities of diverse CCSs densities at different time points of EGF stimulation (Fig. 1e). We compared the differences in densities by quantifying the membrane occupation of all CCSs (Sup. Fig. 1) and individual subclasses (Fig. 1f-h). All CCSs in control cells occupied 0.84±0.2% membrane area and markedly increased 2.5-, 4.6- and 5-fold after EGF stimulation for 5, 15, and 30 min, respectively. The area of all CCSs decreased to near baseline levels after 60 min (0.67±0.33%) (Sup. Fig. 1). Notably, membrane area occupation of FCLs followed a similar time course to all CCSs classes, with a peak at 15 and 30 min post-EGF (3.2±0.58%, 6.5-fold; 3.3±1.92%, 6.7-fold), reaching near-baseline levels at 60 min (0.5±0.22%) (Fig. 1f). In contrast, the mean membrane occupation of domed and spherical clathrin structures remained below 1% with no significant differences across the times tested (Fig. 1g-h). Thus, EGF specifically increased the total area occupied by FCLs at the plasma membrane.

FCLs were considerably bigger after EGF treatment (Fig. 1a-d). The distribution range of the size of hundreds of FCLs changed starting at 5 min EGF treatment (6,753.5-131,167.9 nm^2^), with a peak after 15 min (3,461.5-282,649.1 nm^2^) and 30 min (3,838.7-244,634.4 nm^2^) and approached to control level after 60 min (6,847.3-101,739.3 nm^2^) (Fig. 1i). The average size of domed (21,993.9±4294.8 nm^2^) and spherical clathrin structures (11,075.2±1,714.7 nm^2^) were similar to their respective controls through the time course (Fig. 1j-k). Together, our nanoscale analysis revealed a direct connection between the ultrastructure of FCLs and the temporal response to EGF at the plasma membrane.

### EGFR, Src and β5-integrin are required for flat clathrin lattice formation

EGF triggers phosphorylation cascades that activate distinct cellular effectors. We considered that signaling can regulate FCL biogenesis during growth factor stimulation. To identify the possible members of the EGFR signaling pathway involved in this process, we performed a pharmacological screen (Fig. 2g). We first targeted EGFR using gefitinib (Gefi), a drug that blocks receptor activity by binding to the ATP-pocket in the tyrosine kinase domain^42^. PREM of HSC3 cells preincubated with gefitinib followed by EGF treatment showed a percentage of FCLs membrane area occupation (Gefi+EGF= 0.66 ±0.13%) similar to control cells (Ctrl= 0.48±0.11%) (Fig. 2a-c). Thus, gefitinib decreased the effect of EGF on FCL formation by 4.8-fold (Fig. 2h). We then aimed to evaluate the role of the tyrosine kinase Src for two main reasons.

**Figure 2.**
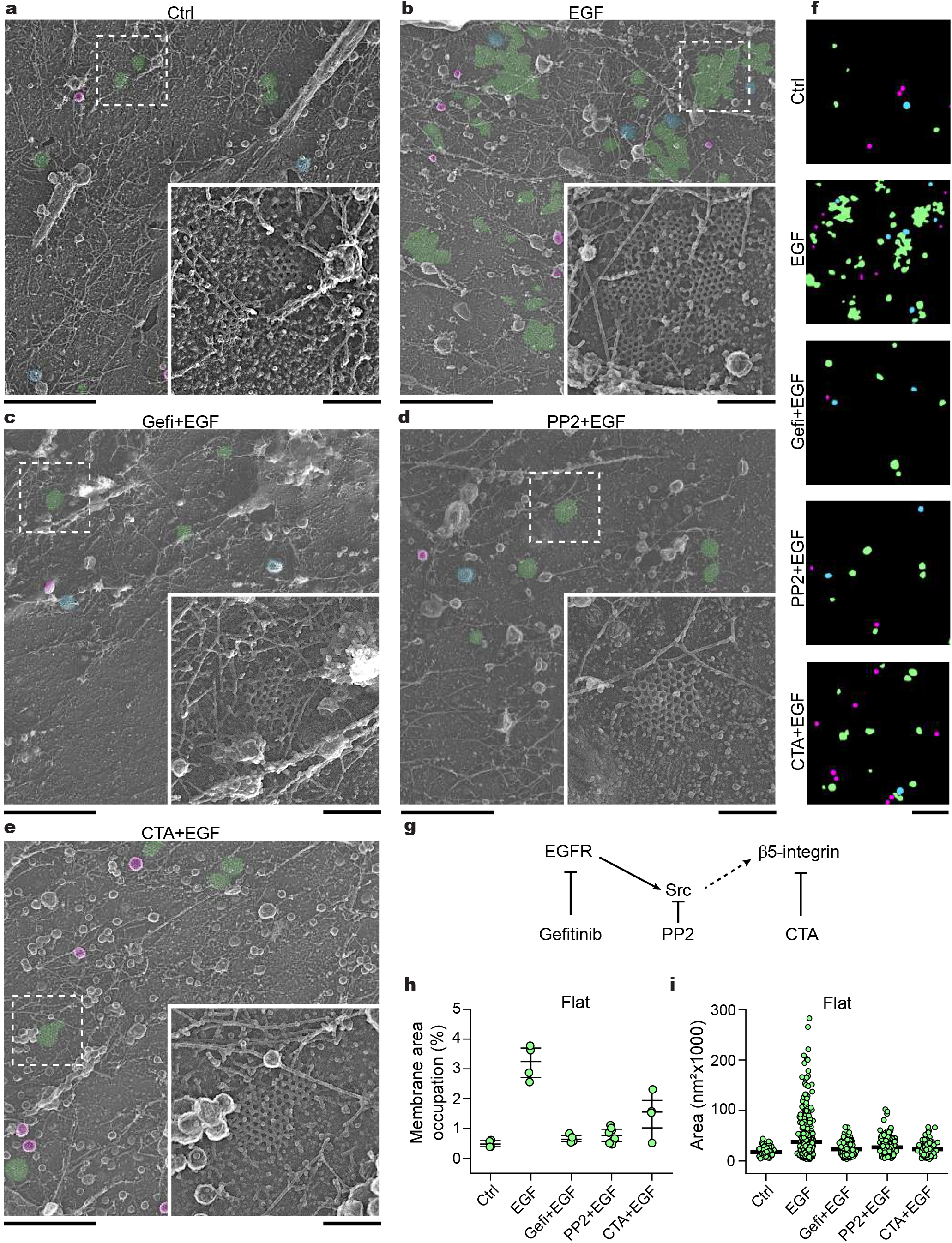
Flat clathrin lattice formation requires EGFR, Src and β5-integrin. **a**, Representative PREMs of control HSC3-EGFR-GFP cells (Ctrl), (**b**) treated either with 50 ng/mL EGF alone (EGF) for 15 min in presence of (**c**) 10 μM gefitinib (Gefi+EGF), (**d**) 10 μM PP2 (PP2+EGF) and (**e**) 10 μM cilengitide acid (CTA+EGF). The magnification insets are shown at the same scale and are outlined with dashed squares in each image. Flat, dome and sphere clathrin-coated structures (CCSs) are shown in green, blue and magenta, respectively, with native grayscale in magnified insets. **f**, Representative masks of segmented cells treated as in (**a-e**). PREM images corresponding to the masks and cropped images are shown in Supplementary Figure 4. **g**, A diagram shows the respective targets of the drugs used in (**a-e**). **h**, Morphometric analysis of the percentage of plasma membrane (PM) area occupation for flat clathrin structures. I-shaped box plots show median extended from 25th to 75th percentiles, and minimum and maximum data point whiskers with a coefficient value of 1.5. **i**, Morphometric analysis of the size of flat structures of cells treated as indicated in (**a-f**). Dot plots show every structure segmented, the bar indicate the median. Ctrl: N_flat_=141, N_cells_=4; EGF: N_flat_=554, N_cells_=4; Gefi+EGF: N_flat_=160, N_cells_=4; PP2+EGF: N_flat_=267, N_cells_=4; CTA+EGF: N_flat_=244, N_cells_=4. Scale bars in (**a-f**) are 1 μm; insets are 200 nm. Ctrl and EGF data are from Figure 1 and shown for reference.

First, Src is a master effector immediately downstream of EGFR. And second, Src has been reported to directly phosphorylate proteins key to the endocytic machinery^43–45^. We used PP2, a specific Src family kinase inhibitor^46^, and then challenged the cells with EGF. We observed a 4.1-fold decrease in FCLs occupation compared to EGF alone, and similar as compared to control cells (PP2+EGF= 0.77 ±0.24%) (Fig. 2d). We also targeted the β5-integrin, a known component of FCLs^19, 34^. We treated cells with cilengitide acid (CTA), a molecule that specifically prevents β5-integrin interaction with the extracellular matrix^47^ followed by EGF stimulation. We detected a 2.2-fold inhibition of FCL stimulation by EGF consistent with the other two drugs tested (CTA+EGF= 1.48. ±0.73%) (Fig. 2e). Representative segmentation masks illustrate the diversity of CCSs in the presence of the three blockers (Fig. 2f). The drugs additionally showed a similar inhibitory effect on the size of FCLs (Fig.2i). The drugs did not significantly affect the percentage of membrane area occupied or the size of domed and spherical clathrin structures, in the presence or absence of EGF (Sup. Fig.3). Overall, these data indicated that activated EGFR, the downstream tyrosine kinase Src, and β5-integrin are all required for the EGF-induced biogenesis of FCLs.

### EGFR, Src and β5-integrin are differentially located in flat clathrin lattices

We envisioned that EGFR, Src and β5-integrin are spatially located in close proximity to FCLs to regulate their formation during EGF signaling. Using total internal reflection fluorescence microscopy (TIRFM) along with a high-throughput correlation analysis^48^, we mapped the location of EGFR, Src and β5-integrin at thousands of individual clathrin-coated structures in an unbiased manner. Correlation analysis illustrates how often proteins of interest are spatially associated with clathrin. We assessed the correlation between EGFR and clathrin. To do this, we used the genome-edited HSC3 endogenously expressing EGFR-GFP and transfected them with mScarlet tagged clathrin light chain A (mScarlet-CLCa). Figure 3a shows clathrin as diffraction-limited puncta, whereas EGFR appears as a more diffuse fluorescent signal across the bottom PM in control cells (C=0.15±0.06). EGF stimulation caused the appearance of pronounced EGFR clusters and a 2.4 - fold increase in its correlation with clathrin (C=0.3±0.15) (see white in overlay). We observed a similar increase in wild type (WT) HSC3 cells co-transfected with EGFR-GFP and mScarlet-CLCa stimulated with EGF (Sup. Fig. 5). We then imaged control cells expressing Src-GFP that showed diffuse florescence similar to EGFR (Fig. 3a). Likewise, the Src signal correlated with clathrin (C=0.41±0.1), but decreased 2.2-fold in the presence of EGF (C=0.18±0.1) (Fig. 3b). Interestingly, the β5-integrin signal appeared as discreet puncta that markedly correlated with clathrin in both control (C=0.79±0.11) and EGF treated cells (C=0.76±0.07) (Fig. 3a-b). Thus, EGFR, Src and, β5-integrin differentially correlate with clathrin. To further confirm this, we measured the EGFR and Src signal using β5-integrin as a reference. Similarly, stimulation increased the β5-integrin correlation with EGFR, but decreased correlation with Src (Sup. Fig. 6). Of note, we tested other members of the β-integrin subfamily including β1, β3, and β6 as well as αV, the most frequent partner of β5-integrin. However, the correlation values were 2-to 4-fold smaller when compared to the β5-integrin correlation with clathrin (Sup. Fig. 7). Altogether, these data support the idea that EGF leads to a persistent recruitment of EGFR and β5-integrin into CCSs with a concurrent loss of Src.

**Figure 3.**
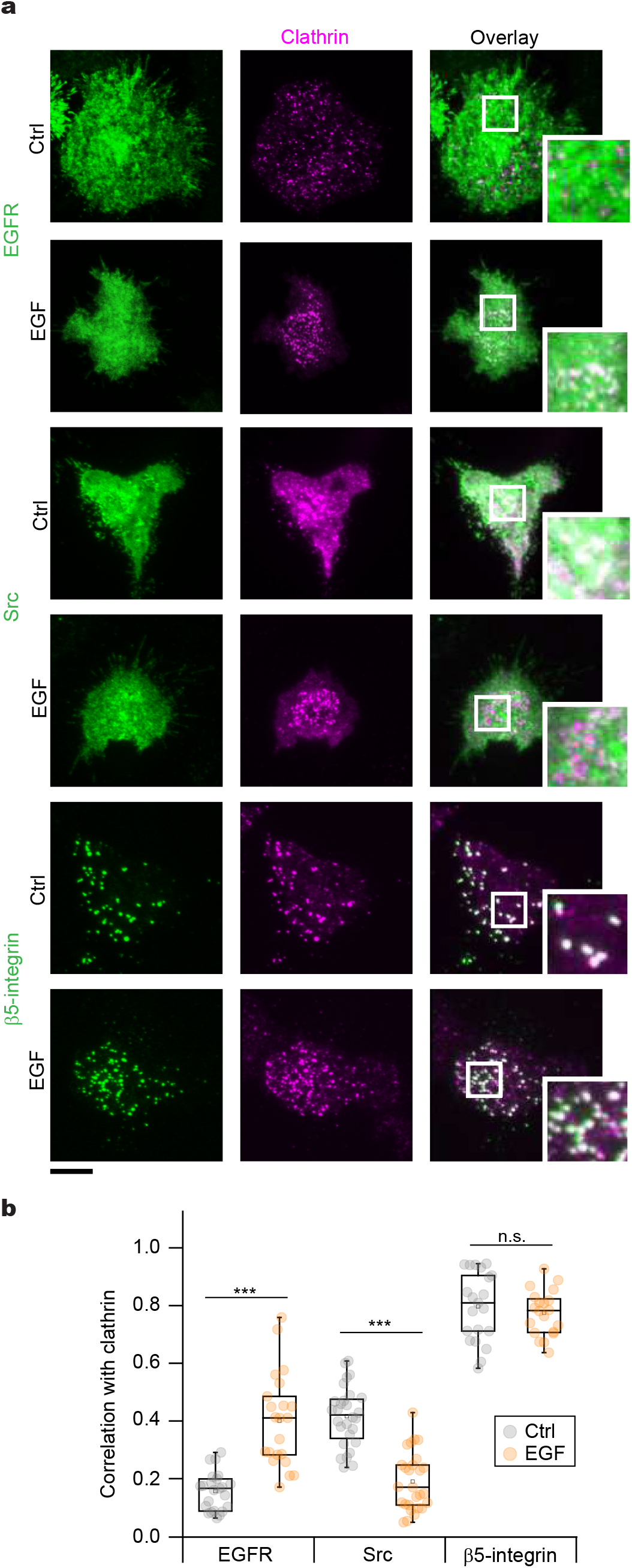
Differential location of EGFR, Src and β5-integrin in clathrin coated structures. **a**, Representative TIRF images of genome-edited HSC3 expressing EGFR-GFP and transfected with mScarlet-CLCa or HSC3 WT cells co-transfected with mScarlet-CLCa + Src-GFP or β5-integrin-GFP before (Ctrl) or after 50 ng/mL EGF stimulation for 15 min. Scale bar is 10 μm; insets are 7.3×7.3 μm. **b**, Automated correlation analysis. Dot box plots show median extended from 25th to 75th percentiles, mean (square) and minimum and maximum data point whiskers with a coefficient value of 1.5. Significance was tested by a two-tailed unpaired t-test. EGFR, \**P=* 5.9×10^−7^; Src, \**P*= 1.7×10^−11^; β5-integrin, ^ns^*P*= 0.358. N_EGFR-Ctrl_= 23 cells – 3728 spots, N_EGFR-EGF_=22 cells – 2173 spots, N_Src-Ctrl_= 28 cells – 1394 spots, N_Src-EGF_= 27 cells –1407spots, N_β5-Ctrl_=21 cells – 1037 spots; N_β5-EGF_=29 cells – 1011 spots.

### EGFR and β5-integrin are connected through Src-mediated phosphorylation

Next we assessed the biochemical connection between EGFR, Src and β5-integrin. EGF binding elicits EGFR dimerization leading to Src tyrosine kinase activation^49^. β5-integrin cytoplasmic domain has three tyrosine residues (Y766, Y774 and Y796) that are highly conserved among their orthologues and paralogues (Sup. Fig. 8). *In silico* analysis suggested Src, among other kinases, has a substantial likelihood of catalyzing phosphorylation at all tyrosines of the β5-integrin cytoplasmic domain (Sup. Fig 8). We therefore hypothesized that these proteins form a signaling loop that can regulate clathrin remodeling through phosphorylation. To support this hypothesis, we used a luciferase-coupled system to measure the phosphorylation of synthetic peptides corresponding to the β5-integrin cytoplasmic domain (Fig. 4a). We detected a 13.6-fold increase in the phosphorylation of the WT peptide in the presence of Src, as compared to the negative control containing only the WT peptide (Fig. 4b). By contrast, this effect was reduced when we tested the non-phosphorylatable peptide 3Y-F. As a positive control, we incubated a peptide corresponding to amino acids 6-20 of p34^cdc2^, a well characterized Src substrate^50^, in the presence of purified and active Src, and we found an incremented phosphorylation. While β5-integrin has been reported to be phosphorylated by PAK4 at S759 and S762^51^, we did not detect phosphorylation when the WT β5-intregrin peptide was incubated with purified PAK4 (Fig. 4b). These data indicate that the intracellular domain of β5-integrin is a bona fide Src substrate *in vitro*.

**Figure 4.**
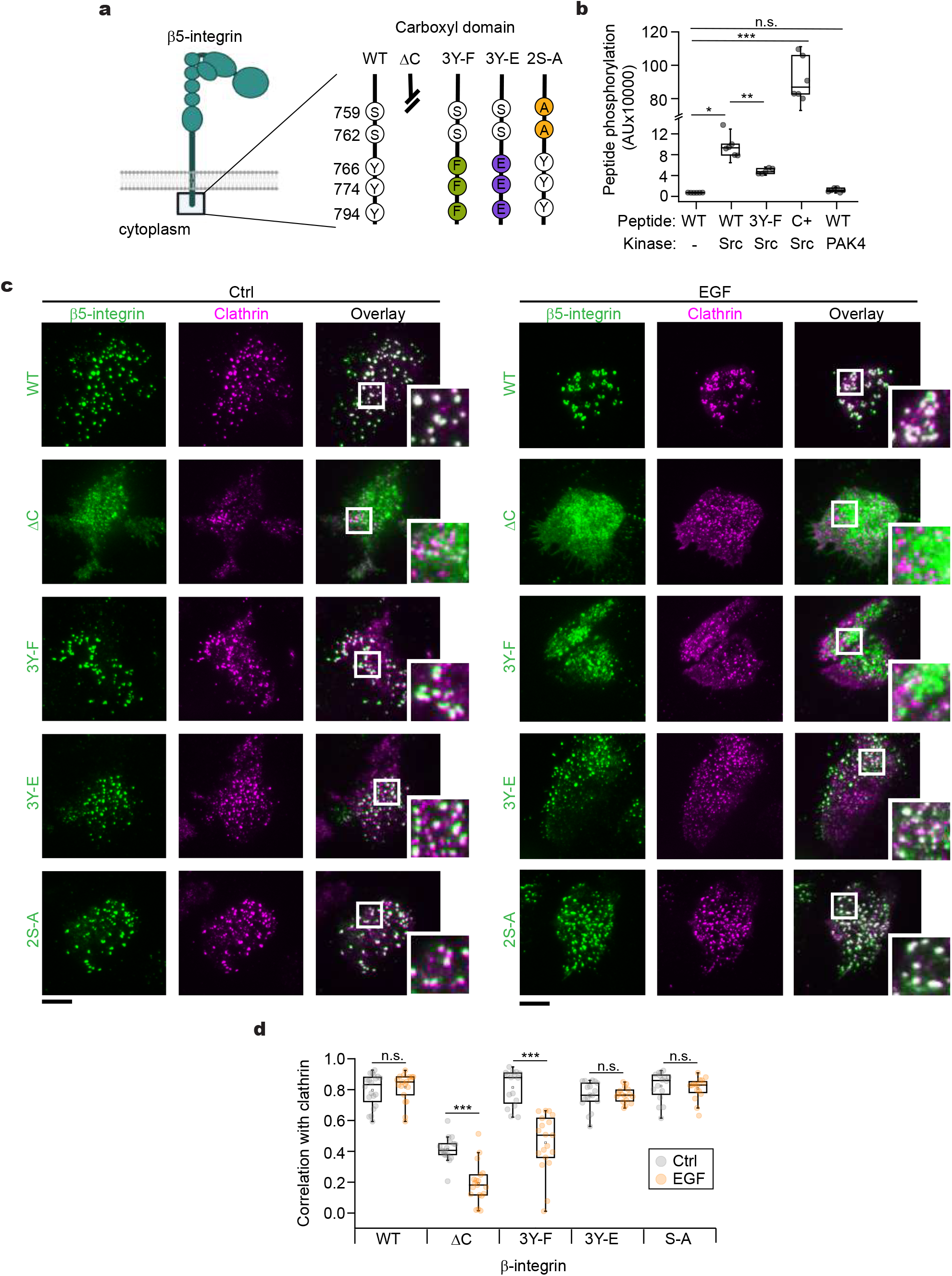
β5-integrin phosphorylation controls spatial correlation with clathrin. **a**, Diagram of β5-integrin and magnification of the cytoplasmic domain showing different mutants. Numbers indicate the residue positions, and letters identify the amino acid. Truncated line in the diagram, indicates deletion of sequence coding for amino acids 743-799, Wild type (WT), carboxyl truncated (ΔC), none-phosphorylatable (3Y-F), phosphomimetic (3Y-E), and PAK-targeted (2S-A). **b**, *In vitro* phosphorylation assay using purified Src or PAK4 and peptides corresponding to the β5-integrin carboxyl domain (742-799) WT and mutants in (**a**). Significance was tested by a two-tailed unpaired t-test, \**P*=1.51×10^−4^, \*\**P*=0.002, \*\*\**P*=1.09×10^−5^, ^ns^P=0.0205. N=4, 6, 5, 6, 6. **c**, Representative TIRF images of HSC3 WT cells co-transfected with mScarlet-CLCa and β5-integrin-GFP WT or containing the different mutations shown in (**a**), either before (Ctrl) or after 50 ng/mL EGF stimulation for 15 min. Scale bars are 10 μm; insets are 7.3×7.3 μm. **d**, Automated correlation analysis of (**a**). Dot box plots show median extended from 25th to 75th percentiles, mean (square) and minimum and maximum data point whiskers with a coefficient value of 1.5. Significance was tested by a two-tailed unpaired t-test, ^ns^*P*_β5-WT_=0.0811, \*\*\**P*_β5-ΔC_=4.03×10^−7^, \*\*\**P*_β5-3Y-F_*=*8.9×10^−5^, ^ns^*P*_β5-3Y-E_*=*0.9149, ^ns^*P*_β5-2S-A_=0.5331. N_β5-WT-Ctrl_=20 cells – 1499 spots, N_β5-WT-EGF_=16 cells – 872 spots, N_β5-ΔC-Ctrl_=17 cells – 1199 spots, N_β5-ΔC-EGF_=19 cells – 1440 spots, N_β5-3Y-F-Ctrl_=15 cells – 826 spots, N_β5-3Y-F-EGF_=17 cells – 1099 spots, N_β5-3Y-E-Ctrl_=17cells – 1245 spots, N_β5-3Y-E-EGF_=16 cells – 1122 spots, N_β5-2S-A-Ctrl_=16 cells – 1326 spots, N_β5-2S-A-EGF_=16 cells – 953 spots.

To evaluate the cellular effects of tyrosine phosphorylation of β5-integrin, we co-transfected WT HSC3 cells with mScarlet-CLCa plus either the β5-intregrin-GFP wild type (WT), carboxyl-truncated (ΔC), non-phosphorylatable (3Y-F), phosphomimetic (3Y-E), or PAK targeted (2S-A) mutants (Fig. 4a). We tested this mutants using two-color TIRFM as previously described. In control cells, we observed that the fluorescent signal of either WT, 3Y-F, 3Y-E or 2S-A mutants appeared as diffraction-limited punctate that highly correlated with clathrin (C=0.79±0.1, 0.81±0.11, 0.76±0.08, 0.82±0.09, respectively) (Fig. 4c-d). In contrast, the ΔC mutant exhibited a diffuse fluorescence across the membrane and a greatly decreased correlation with clathrin (C=0.41±0.08). In cells stimulated with EGF, we observed that the WT β5-integrin strongly correlated with clathrin (C=0.84±0.05) (Fig. 4 and Fig. 5c-d). This spatial correlation was abolished in the ΔC (C=0.18±0.12) and the 3Y-F mutant (C=0.46±0.21). By contrast, the correlation with clathrin persisted in the 3Y-E (C=0.76±0.04) and the 2S-A mutants (C=0.8±0.07). Thus, β5-integrin requires tyrosine phosphorylation to spatially correlate with clathrin. Moreover, these experiments suggest that crosstalk between EGFR and β5-integrin mediated by Src takes place at CCSs.

**Figure 5.**
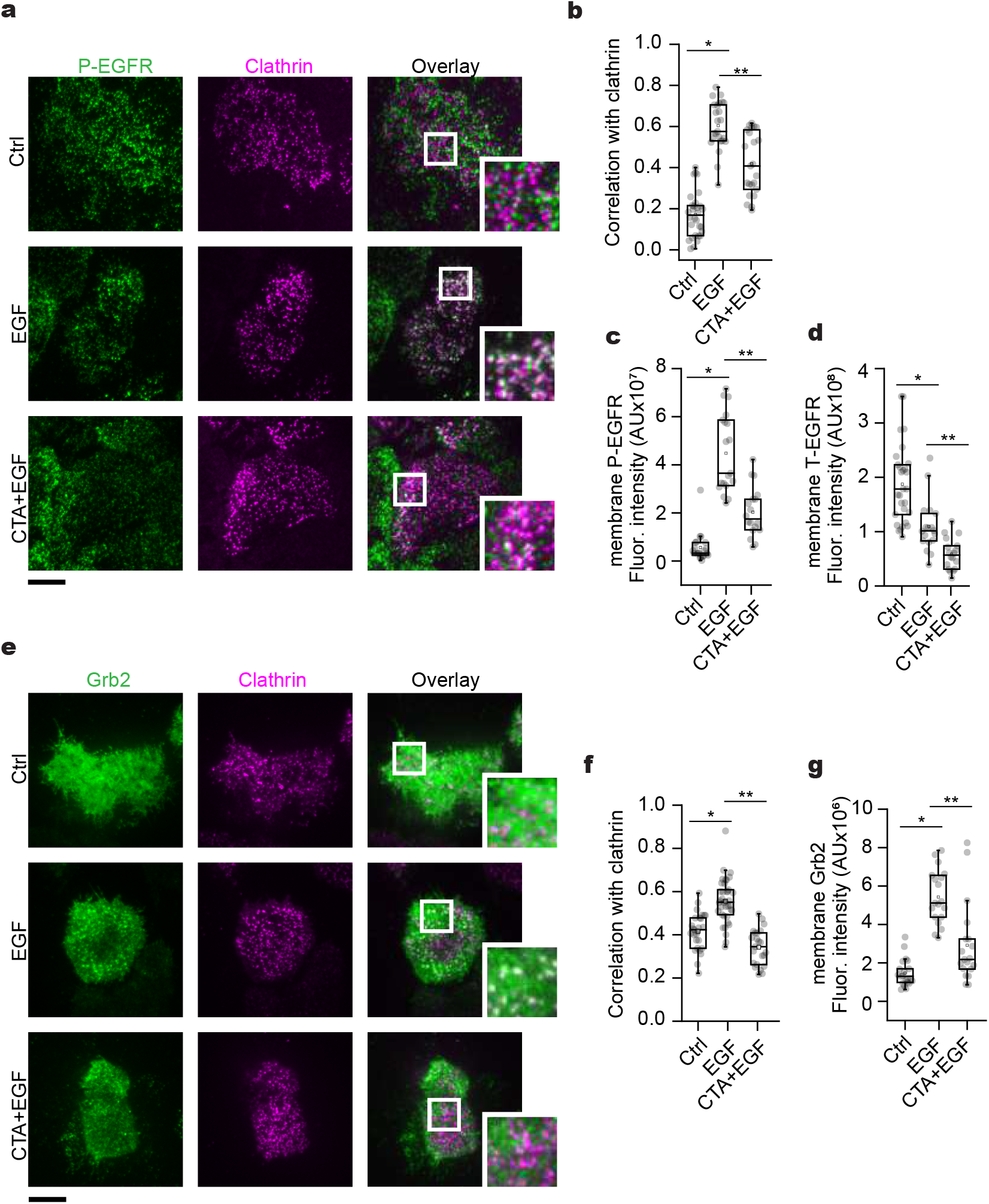
Flat clathrin lattices partition sustained signals at the plasma membrane. **a**, Representative TIRF images of control (Ctrl) unroofed genome-edited HSC3 cells expressing EGFR-GFP transfected with mScarlet-CLCa and immunolabeled with anti-phospho EGFR (P-EGFR) coupled to Alexa 647, treated with 50 ng/mL EGF alone (EGF) or in the presence of 10 μM cilengitide acid (CTA+EGF). **b**, Automated correlation analysis of (**a**). \**P*=1.11×10^−19^, \*\**P*=1.70×10^−5^. N_Ctrl_=30 cells, N_EGF_=25 cells, N_CTA+EGF_=23 cells. **c**, Fluorescence intensity measurements of the signal from P-EGFR. \**P*=1.13×10^−9^, \*\**P*=4.33×10^−6^. N_Ctrl_=21 cells, N_EGF_=19 cells, N_CTA+EGF_=18 cells. **d**, Fluorescence intensity measurements of the signal from total EGFR-GFP (T-EGFR). \**P*=1.95×10^−4^, \*\**P*=6.00×10^−4^. N_Ctrl_=29 cells, N_EGF_=18 cells, N_CTA+EGF_=18 cells. **e**, Representative TIRF images of HSC3 WT cells transfected with mScarlet-CLCa and Grb2-GFP before (Ctrl) or after 15 min EGF stimulation. **f**, Automated correlation analysis of (**e**). \**P*=2.43×10^−7^, \*\**P*=1.06×10^−11^. N_Ctrl_=27 cells, N_EGF_=38 cells, N_CTA+EGF_=23 cells. **g**, Fluorescence intensity measurements of the signal coming from immunolabeled Grb2. \**P*=2.17×10^−11^, \*\**P*=1.06×10^−4^. N_Ctrl_=21 cells, N_EGF_=19 cells, N_CTA+EGF_=19 cells. Scale bars are 10 μm; insets are 7.3×7.3 μm square. Dot box plots show median extended from 25th to 75th percentiles, mean (square) and minimum and maximum data point whiskers with a coefficient value of 1.5. Significance was tested by a two-tailed unpaired t-test. AU, fluorescence arbitrary units.

### Flat clathrin lattices regulate sustained signaling at the plasma membrane

Finally, we hypothesized that FCLs mediate EGFR membrane-associated signal transduction by regulating the distribution of active receptors and downstream interactors in both space and time. Using TIRFM, we visualized the presence of active EGFR at the PM by immunolabeling endogenous phosphor-Y1068 (P-EGFR), a well-established marker of EGFR activity (Fig. 5a). In stimulated cells, the correlation of P-EGFR with clathrin increased 3.5-fold (C=0.6 ±0.11) with respect to the control (C=0.17 ±0.1). Conversely, disruption of FCLs formation by CTA treatment (Fig.2e), decreased the P-EGFR correlation with clathrin by ~30% (C=0.41 ±0.14) (Fig 5b). A similar trend was observed when we measured the fluorescence intensity of P-EGFR at the PM (Fig. 5c). Furthermore, we measured the fluorescence intensity of the total EGFR-GFP signal (T-EGFR) at the PM as an indicator of receptor internalization. We observed that EGF led to a 41.7% decrease of the T-EGFR, and CTA treatment further decreased receptor at the PM (68.7%) (Fig. 5d). We also examined the location of the downstream master scaffold Grb2 in FCLs (Fig. 5e). While EGF caused an increase in both Grb2 correlation with clathrin and recruitment of the adaptor to the PM, CTA blocked these effects (Fig. 5e–). Together these data suggest that FCLs activate growth factor signals at the PM by delaying endocytosis of a population of active EGFR along with key partner proteins such as Grb2. These events are mediated by the phosphorylation-dependent cross talk between EGFR, integrins, and clathrin at the plasma membrane.

## DISCUSSION

Crosstalk between signaling systems allows for biological processes to be integrated, responsive, and adaptable^52^. There is an emerging hypothesis that FCLs can act as signaling zones or adhesion sites at the PM, filling unique roles outside of endocytosis. Here, we show that activated EGFR, Src, and β5-integrin are coupled to a dramatic growth and maintenance of FCLs in human cells. These planar clathrin sites in turn partition and enhance growth factor signals at the PM (Fig. 6). Thus, two receptor systems (growth factor and adhesion) are connected, clustered, and controlled at the nanoscale by endocytic proteins. We propose that a reciprocal feedback loop operates where FCLs facilitate local crosstalk between EGFR, β5-integrin, and other signaling proteins to create dynamic signaling hubs across the PM.

**Figure 6.**
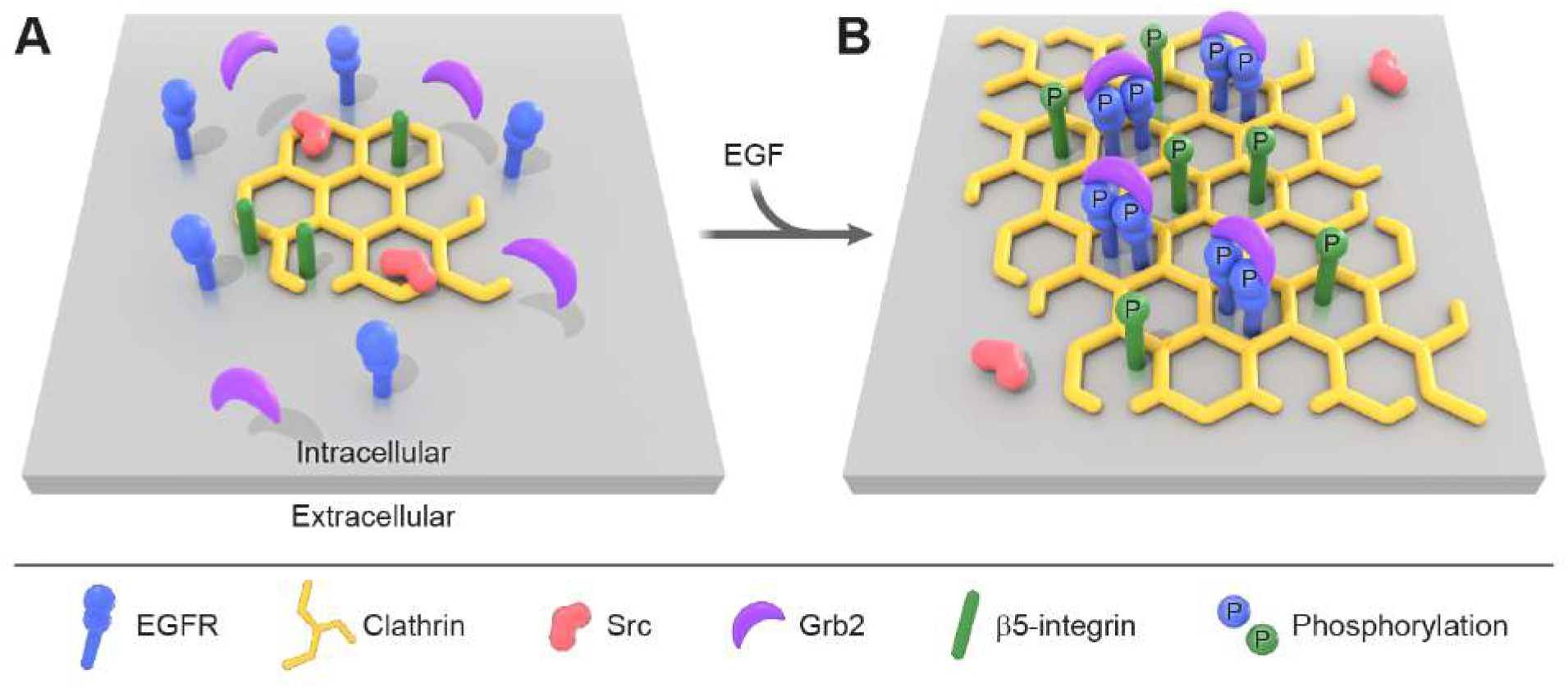
Model of flat clathrin lattices biogenesis during growth factor response. The critical steps of the model are: 1) small flat clathrin lattices are in close proximity to Src and are highly enriched with β5-integrin; 2) EGF triggers the dimerization, clustering and cross-phosphorylation of EGFR at growing FCLs; 3) this in parallel allows the biding of the downstream scaffold Grb2 and locally activates Src; 4) which in turn phosphorylates β5-integrin cytoplasmic domain; 5) the maintenance of the EGFR/Src/β5-integrin axis promotes flat clathrin lattice growth. A key implication of this model is that two different receptor systems are spatially organized at the nanoscale within flat clathrin lattices.

First, we observed clathrin using platinum replica electron microscopy to structurally distinguish flat from curved clathrin structures. Surprisingly, EGFR activation resulted in a dramatic increase in FCLs size and density. Other shapes of clathrin were unchanged. Blocking the receptor abolished these effects. Of note, the structural changes we see follow a time course similar to the activation dynamics of downstream kinases^53^, further supporting the direct connection between clathrin remodeling and cell signaling. Our pharmacological screening linked the tyrosine kinase Src and the adhesion receptor β5-integrin to FCL formation and EGFR activation. Historically, growth factor receptors and integrins have been biochemically connected to Src in several ways^10^. Here, we show a new direct spatial connection. We also found that FCLs are preloaded with a subpopulation of Src. This kinase was released from the complex in response to EGF. In contrast, β5-integrin is enriched in FCLs^19, 34, 35^, and we found that this correlation with clathrin persisted in response to EGF. Thus, EGFR, Src and β5-integrin are dynamically coupled through FCLs to regulate EGF signaling.

We showed that this new pathway is controlled by phosphorylation. Specifically, *in silico* analysis and biochemical assays indicated that the β5-integrin intracellular domain is a Src substrate. Src activation kinetics are fast and occur within 5 minutes^54^. Thus, it is possible that at an early stage of clathrin growth, Src phosphorylates targets and is then released. While our results point toward an early phosphorylation event in β5-integrin mediated by Src, it is also possible that Src continually cycles on and off clathrin during receptor activation and plays a more extensive role in the process. Likewise, Src is a promiscuous kinase and might phosphorylate additional substrates located on other PM structures such as caveolae and focal adhesions^55, 56^.

Second, we found that deletion of the β5-integrin cytoplasmic domain and non-phosphorylatable mutations (3Y-F) block integrin association with clathrin. These effects were rescued by a phosphomimic mutant (3Y-E), suggesting that tyrosine phosphorylation of β5-integrin is a molecular switch in this process. Interestingly, the β5-integrin cytoplasmic domain interacts with endocytic proteins including Eps15, ARH, and Numb^19^. Integrin cytoplasmic tails can also induce profound differences in the behavior of integrins^57^. Thus, we propose that a phosphorylation switch in β5-integrin is the regulator for the orchestrated recruitment of the endocytic machinery to sites where FCLs form and grow. In this regard, the growth factor response is directly linked to the cellular adhesion system through activation of endocytic proteins and controlled by phosphorylation.

A recent hypothesis proposes that long-lived FCLs arise from adhesive forces generated from integrins that physically prevent clathrin from curving—a process called frustrated endocytosis^34 27, 58^. Interestingly, we observed the FCL formation and the enrichment of active EGFR and Grb2 peak after 15 min of stimulation with EGF (Fig. 1 and 5). At the same time, we detected a decrease in the overall signal of EGFR at the PM. This decrease suggests that EGF triggers the internalization of a population of EGFR into the cell. In parallel, some phosphorylated and active receptors remain at the PM in clathrin. By preventing endocytosis of a subpopulation of EGFR, this clathrin/adhesion complex could prolong EGF signaling at these sites. These domains might also act as diffusion traps for EGFR and other growth factor receptors whose diffusion decreases after agonist stimulation^59–62^. Using fluorescence microscopy, EGFR and other structurally diverse receptors have been reported to form long-lived complexes^27, 63–66^. In contrast, stimulation of LPA1 receptor has been shown to trigger the depolymerization of FCLs^33^. Thus, different systems might activate or deactivate these structures to regulate their activity. How endocytosis, adhesion, and receptor diffusion cooperate across the entire population of active receptors to control signaling will be an important future area of study.

Is this mechanism unique to EGFR? Many receptors including 7-transmembrane receptors and B cell receptors trigger clathrin nucleation at the PM upon biding their ligands^27, 63–66^. For the β_2_-adrenergic receptor, the increase in clathrin occurs with a delay in clathrin-coated vesicle maturation and no differences in the overall rate of vesicle scission^64^. Our data revealed a similar increase in clathrin nucleation during EGF stimulation, but the major changes to clathrin occurred specifically and exclusively with a dramatic growth in FCLs. It is possible that other receptor cargos also stabilize flat clathrin coats to act as nanoscale receptor signaling domains. Thus, FCLs could be generalized signaling hubs at the PM. Future work is needed to test this hypothesis.

Growing evidence suggests that signaling systems are locally organized by organelles and cytoskeletal structures. We propose that FCLs are a unique plasma membrane scaffold that dynamically capture and organize receptors and signaling molecules in space and time through multivalent interactions at the nanoscale. We suggest that crosstalk between EGFR and β5-integrin through Src phosphorylation occurs in FCLs and simultaneously regulates their biogenesis. Finally, because of the importance of EGFR/Src/β5-integrin in physiology, and the connection between dysregulation of this system and cancer, FCLs likely play a broader role in coordinating the cellular responses to chemical and mechanical stimuli. Understanding these pathways is key to understanding cellular functions in both health and disease. Our data provide a new nanoscale signaling platform that dynamically organizes, coordinates, and regulates this essential biology.

## METHODS

### Cell culture and transfection

Wild-type HSC-3 (human oral squamous carcinoma) cells were obtained from the JCRB Cell Bank (JCRB0623). Genome-edited HSC-3 cells expressing endogenous EGFR-GFP were previously reported^39^ and kindly donated by Dr. Alexander Sorkin (University of Pittsburgh). Cells were grown at 37 °C with 5% CO_2_ in phenol-free Dulbecco’s modified Eagle’s medium (DMEM) (Thermo-Fisher, Gibco™, 31053028) containing 4.5 g/L glucose and supplemented with 10% (v/v) fetal bovine serum (Atlanta Biologicals, S10350), 50 mg/mL streptomycin - 50 U/mL penicillin (Thermo-Fisher, Gibco™, 15070063), 1% v/v Glutamax (Thermo-Fisher, 35050061), and 1 mM sodium pyruvate (Thermo-Fisher, Gibco™, 11360070). Cell lines were used from low-passage frozen stocks and monitored for mycoplasma contamination. For experiments, cells were grown on 25 mm diameter rat tail collagen I-coated coverslips (Neuvitro Corporation, GG-25-1.5-collagen). For transfections, cells were incubated for 4 h with 500 ng of the indicated plasmid(s) and 5 μL of Lipofectamine 3000 (Thermo-Fisher, L3000015) in OptiMEM (Thermo-Fisher, Gibco™, 31985062). Experiments were performed 18 h after transfection.

### Plasmids

EGFR-GFP #32751, Src-GFP #110496, Src-mCherry #55002, αV-integrin-mEmerald #53985, β1-integrin-GFP #69804 were purchased from Addgene. β5-integrin-GFP was kindly donated by Dr. Staffan Strömblad (Karolinska Institutet). EGFR-mScarlet, mScarlet-CLCa, β3-integrin-GFP, β6-integrin-GFP, β5-integrin-GFP lacking 743-799 amino acids (ΔC), β5-integrin-GFP containing point mutations Tyr766, 774, 794Phe (3Y-F), Tyr766, 774, 794Glu (3YE), and Ser759, 762Ala (2S-A), were built using either Q5 Site-Directed Mutagenesis Kit (New England Biolabs, E0554S) or In-Fusion HD Cloning Plus (Clonetech, 638911) following manufacturer’s instructions. All plasmids were confirmed by sequencing (Psomagen).

### EGF pulse-chase stimulation and drug treatments

Cells were incubated in starvation buffer (DMEM containing 4.5 g/L D-glucose, supplemented with 1% v/v Glutamax and 10 mM HEPES) for 2 h before the pulse-chase assay. Then, cells were pulsed in starvation buffer supplemented with 0.1% w/v bovine serum albumin at 4 °C for 40 min with 50 ng/mL human recombinant EGF (Thermo-Fisher, Gibco™, PHG0311L) to allow ligand bind to the EGFR. In brief, cells were washed twice with PBS (Thermo-Fisher, Gibco™, 10010023). Synchronized receptor activation and endocytosis were triggered by placing the coverslips in pre-warmed media and incubation at 37 °C for the indicated times. To stop stimulation, cells were washed twice with iced-cold PBS. To block EGFR, Src, and β5-integrin cells were incubated for 20 min before chase and during pulse with 10 μM gefitinib (Santa Cruz Biotechnology, 184475-35-2), 10 μM PP2 (Thermo-Scientific, 172889-27-9), and 10 μM cilengitide acid (CTA) (Sigma-Aldrich, SML1594), respectively.

### Cell unroofing and fixation

After EGF pulse-chase stimulations, cells were rinsed briefly with stabilization buffer (30 mM HEPES, 70 mM KCl, 5 mM MgCl_2_, 3 mM EGTA, pH 7.4). Fixed cell membranes were obtained with application of unroofing buffer containing 2 % paraformaldehyde in stabilization buffer using a 10 mL syringe with a 22 gauge, 1.5 needle. The syringe was held vertically within 1 cm of the coverslip during the mechanical unroofing. Afterwards, the coverslips were moved to fresh unroofing buffer containing 2 % paraformaldehyde for 20 min. They were rinsed 4× with PBS followed by electron or fluorescent microscopy preparation.

### Platinum replica electron microscopy (PREM)

Coverslips were transferred from glutaraldehyde into 0.1% w/v tannic acid for 20 min. Then, coverslips were rinsed 4× with water, and placed in 0.1% w/v uranyl acetate for 20 min. The coverslips were dehydrated, critical point dried, and coated with platinum and carbon as previously described^40^. The replicas were separated from glass coverslips using hydrofluoric acid and mounted on glow-discharged Formvar/carbon-coated 75-mesh copper TEM grids (Ted Pella 01802-F).

Transmission Electron Microscopy imaging was performed as previously described ^67^ at 15,000× magnification (1.2 nm per pixel) using a JEOL1400 (JEOL) and SerialEM software for montaging. Montages were stitched together using IMOD^67^. Images are presented in inverted contrast. Each montage was manually segmented in imageJ^68^ by outlining the edge of the membrane, flat clathrin structures (no visible curvature), domed clathrin structures (curved but can still see the edge of the lattice), and sphere clathrin structures (curved beyond a hemisphere such that the edge of the lattice is no longer visible) as previously described^41^. The percentage of occupied membrane area was defined as the sum of areas from clathrin of the specified subtype divided by the total area of visible membrane.

### Immunofluorescence

Unroofed cells were incubated in PBS containing 3% w/v bovine serum albumin (Fisher Bioreagents, BP9703) and 0.1% v/v triton X-100 (Sigma-Aldrich, T9284) for 1 h at room temperature. The cells were then immunolabelled with the indicated primary antibodies for 1 h at room temperature: 1:1000 anti-Clathrin Heavy Chain monoclonal antibody X22 (Thermo-Fisher, MA1-065), 1:800 anti-Phospho-EGF Receptor (Tyr1068) (D7A5) XP^®^ Rabbit mAb (Cell Signaling, 3777), 1:50 anti-Grb2 Y237 (Abcam, 32037). Then, cells were washed 5x and incubated in 2.5 μg/mL of the corresponding secondary antibody conjugated with Alexa Fluor 647 for 30 min at room temperature (Invitrogen, anti-mouse A21237, anti-rabbit A21246). When indicated, cells were labeled with 16.5 pmol of Alexa Fluor 488-Phalloidin for 15 min (Invitrogen, A12379). The coverslips were then rinsed 4× with blocking buffer, 4× with PBS, and then post-fixed with 4% paraformaldehyde in PBS for 20 min and imaged immediately or refrigerated overnight.

### Total Internal Reflection Microscopy (TIRFM)

Cells were imaged on an inverted fluorescent microscope (IX-81, Olympus), equipped with a 100x, 1.45 NA objective (Olympus). Combined green (488 nm) and red (561 nm) lasers (Melles Griot) were controlled with an acousto-optic tunable filter (Andor) and passed through a LF405/488/561/635 dichroic mirror. Emitted light was filtered using a 565 DCXR dichroic mirror on the image splitter (Photometrics), passed through 525Q/50 and 605Q/55 filters and projected onto the chip of an electron-multiplying charge-coupled device (EMCCD) camera. Images were acquired using the Andor IQ2 software. Cells were excited with alternate green and red excitation light, and images in each channel were acquired at 500-ms exposure at 5 Hz. Automated correlation analysis was performed on aligned images as described previously^48^ and the fluorescence intensity signal at the plasma membrane was assessed using ImageJ software by measuring the integrated density (mean gray value) of the background subtracted from that of the cell and normalizing this value to the total cell area.

### *In silico* analysis

Integrin sequences were obtained from the UniProt Knowledgebase. β5-integrin orthologs: *H. sapiens* (ID P18084); *M. musculus* (ID O70309); *B. taurus* (ID P80747); *P. cynocephalus* (ID Q07441); *X. laevis* (ID Q6DF97); *D. rerio* (ID F1Q7R1). Integrin orthologs: β1-integrin (ID P05556); β2-integrin (ID P05107); β3-integrin (ID P05106); β6-integrin (ID P18564); β7-integrin (ID P26010). The sequence alignments were performed using the blast-protein suite (protein-protein BLAST, (http://www.uniprot.org/blast/). The prediction of phosphorylation sites was obtained using NetPhos 3.0 (http://www.cbs.dtu.dk/services/NetPhos) and the Group-based Prediction System, GPS 2.0 (http://gps.biocuckoo.org/) on-line services, employing cut-off values of 0.75 and 4, respectively. Prediction of the probable protein kinases involved was obtained using GPS 2.0.

### *In vitro* kinase assay

Identification of the β5-integrin carboxyl domain as Src substrate was determined using the ADP-Glo Kinase Assay (Promega, V6930) following protocols recommended by the manufacturer. All reactions were performed in kinase buffer (40 mM Tris, pH 7.5, 20 mM MgCl_2_, 2 mM MsSO_4_, 100 mM Na_3_VO_4_, 10 mM DTT) supplemented with 50 mM ATP, 1 mM of the indicated peptide, and 50 ng of purified active Src (Millipore-Sigma, 14-326) or PAK (Millipore-Sigma, 14-584). β5-integrin peptide corresponding to amino acids 743-799 and Y766, 774, 794F were chemically synthetized (Biobasic). As a positive control we used a bona fide Src substrate peptide corresponding to amino acids 6-20 of p34^cdc2^ (Millipore-Sigma, 12-140). The reactions were carried out at room temperature in a total volume of 25 μL for 40 min in white 96-well, F-bottom, non-binding microplates (Greiner Bio-one, 655904). Signal was recorded using a luminometer (Biotek) with an integration time of 0.5 s.

### Statistics

Data were tested for normality and equal variances with Shapiro–Wilk. The statistical tests were chosen as follows: unpaired normally distributed data were tested with a two-tailed *t*-test (in the case of similar variances) or with a two-tailed *t*-test with Welch’s correction (in the case of different variances). All tests were performed with Origin 2015.

## Data availability

All data supporting this work are available upon request to the corresponding author.

## Author contributions

MAAM, KAS and JWT designed experiments. KAS developed software for data analysis. MAAM performed experiments and analyzed data. MPS performed molecular cloning and helped with in vitro phosphorylation assays. MAAM wrote and JWT edited the manuscript and all authors commented on the work. JWT supervised the project.

## Acknowledgements

We would like to thank the NHLBI Electron Microscopy core for support with EM imaging and instrumentation, Xufeng Wu and the NHLBI light Microscopy core for support with fluorescence imaging and instrumentation, Ethan Tyler of NIH Medical Arts Department for the creation of Figure 6, Agila Somasundaram, and members of the Taraska laboratory for discussion and comments on the manuscript. JWT is supported by the NHLBI Intramural Research Program, National Institutes of Health, Bethesda, Maryland.

## SUPPLEMENTARY FIGURE LEGENDS

**Supplementary Figure 1.**
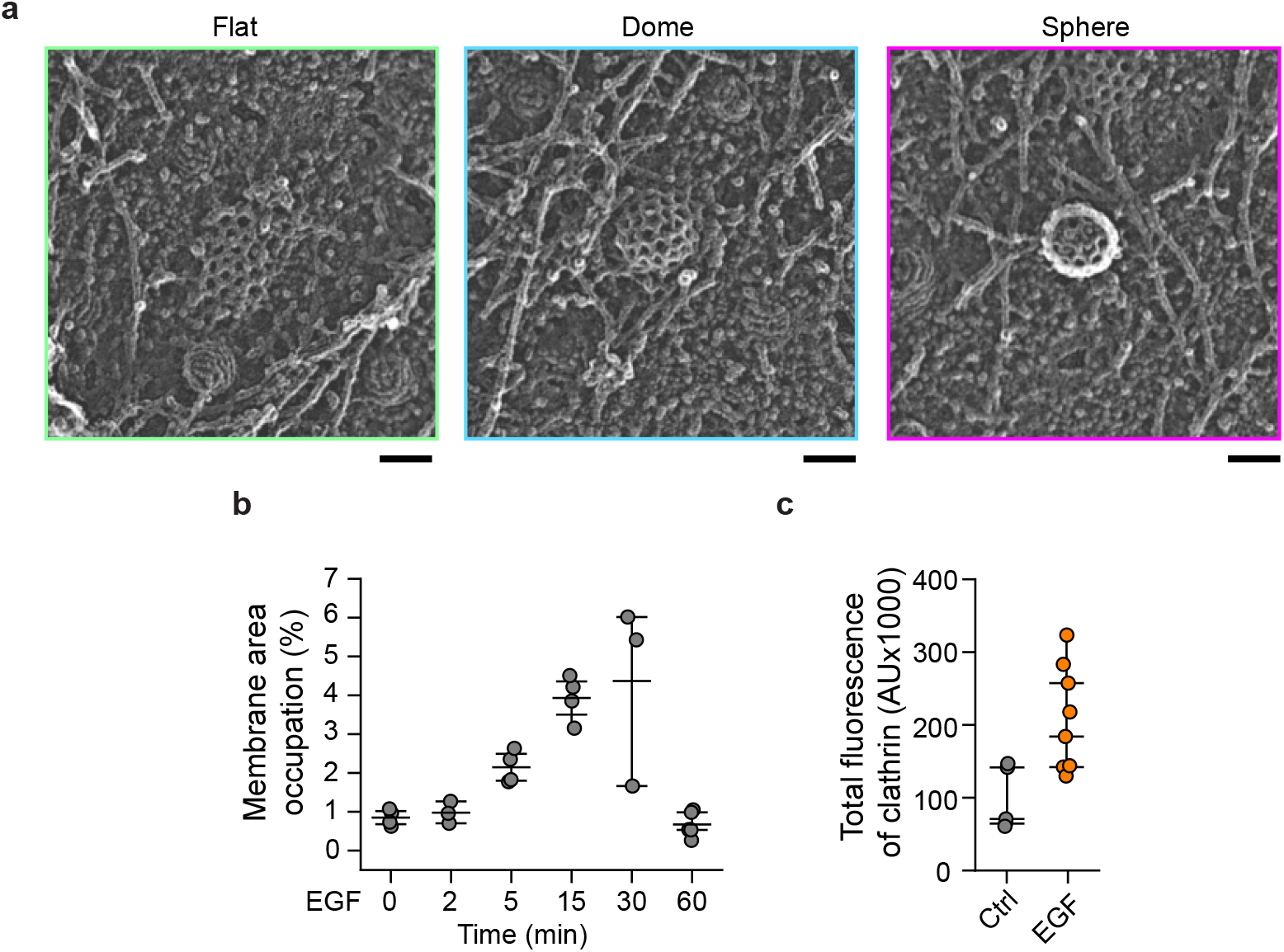
EGF increases the density of clathrin at the plasma membrane. **a**, Representative PREM images of the flat (green), dome (blue), and sphere (magenta) clathrin-coated structures (CCSs) segmented in Fig. 1. Scale bars are 100 nm. **b**, Morphometric analysis of the percentage of plasma membrane area occupation for all clathrin-coated structures (CCSs) in PREM images of control (Ctrl) HSC3-EGFR-GFP cells or treated with 50 ng/mL EGF for 2, 5, 15, 30 and 60 min from Figure 1. I-shaped box plots show median extended from 25th to 75th percentiles, and minimum and maximum data point whiskers with a coefficient value of 1.5. 0 min: N_cells_=4; 2 min: N_cells_=3; 5 min: N_cells_=4; 15 min: N_cells_=4; 30 min: N_cells_=3; 60 min: N_cells_=5. **c**, Fluorescence intensity measurements of the signal from clathrin heavy chain. Control (Ctrl) unroofed HSC3-EGFR-GFP cells or treated with 50 ng/mL EGF were immunolabeled with anti-clathrin heavy chain coupled to Alexa 647. Ctrl: N_cells_=5; EGF: N_cells_=8.

**Supplementary Figure 2.**
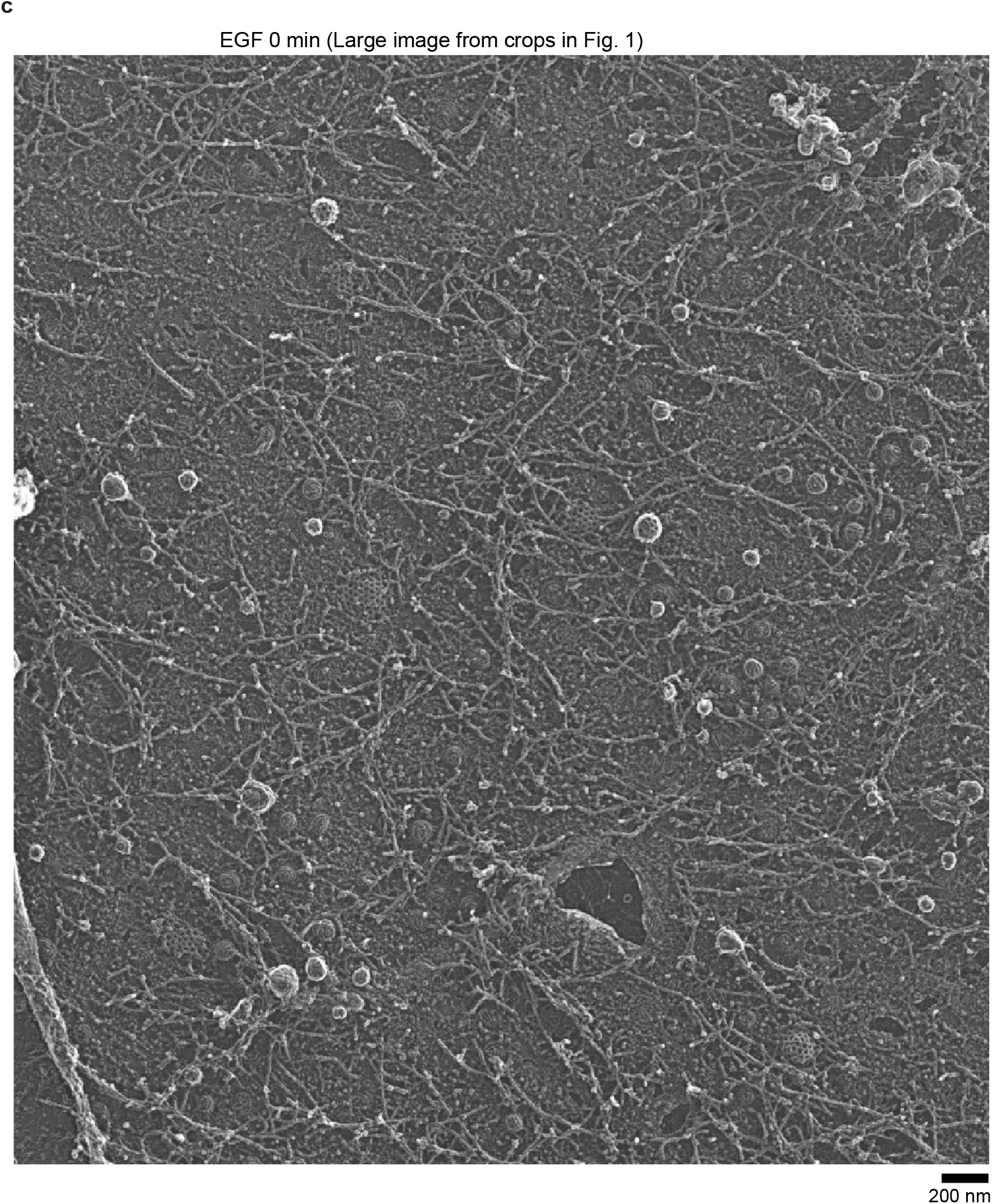
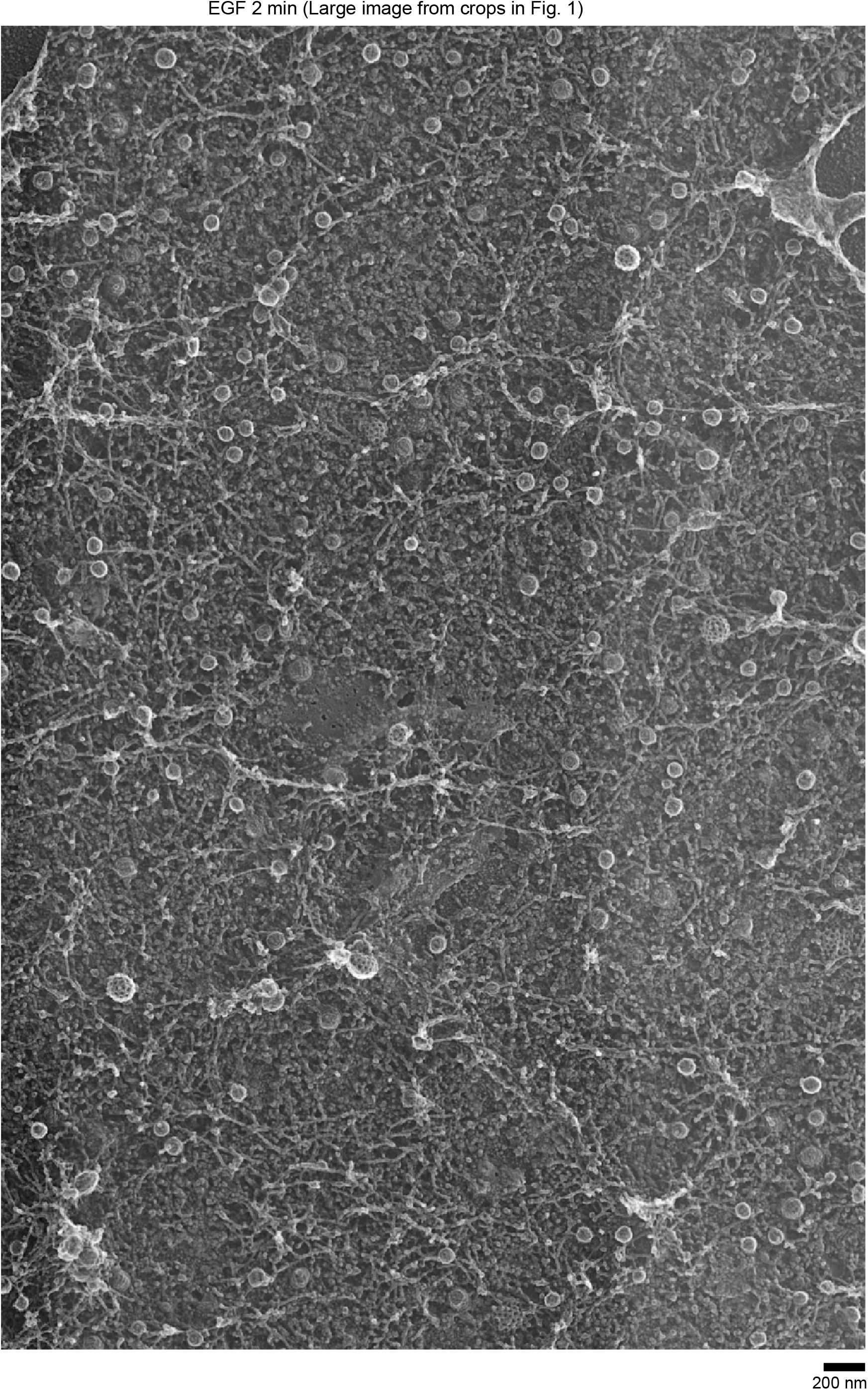
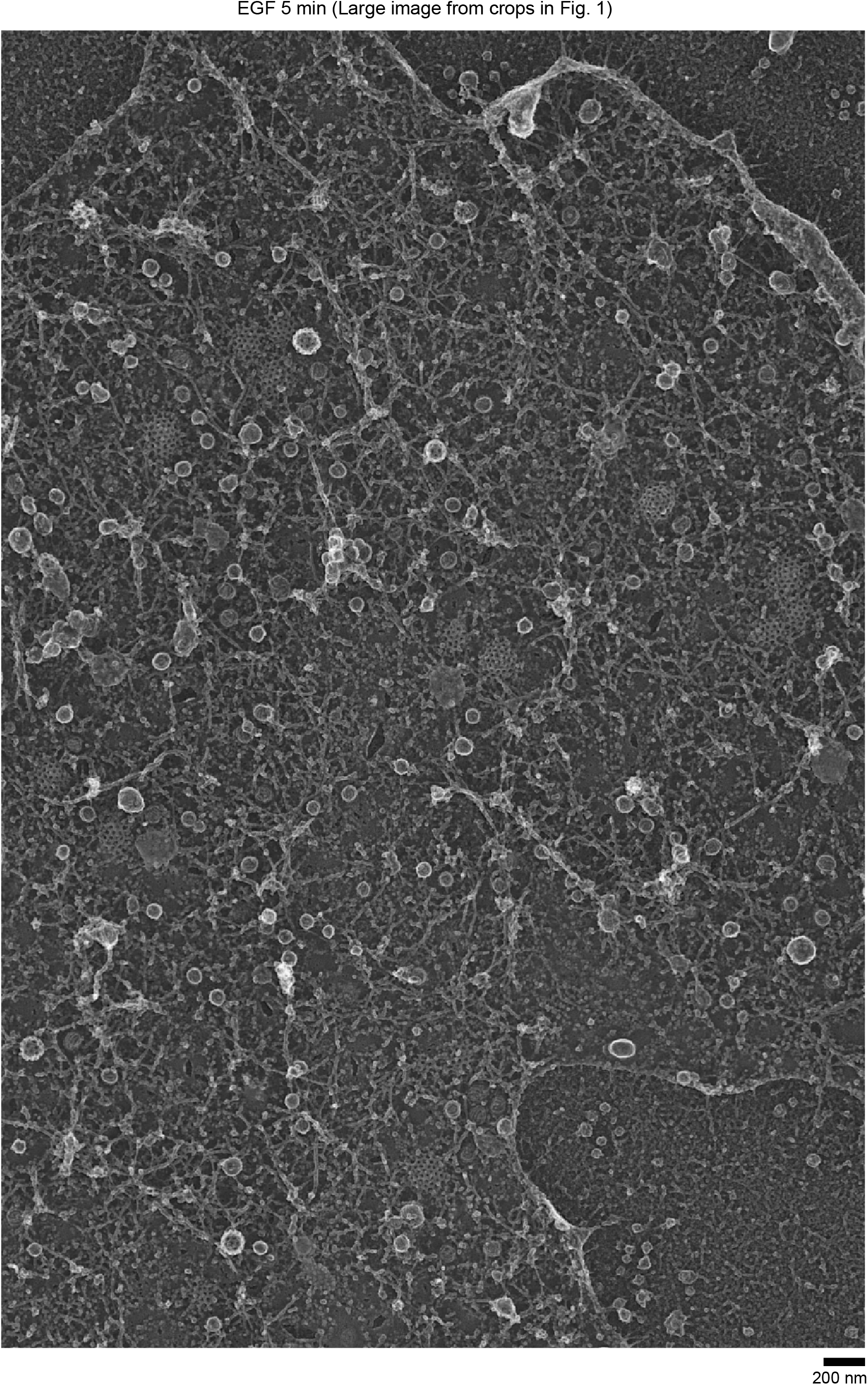
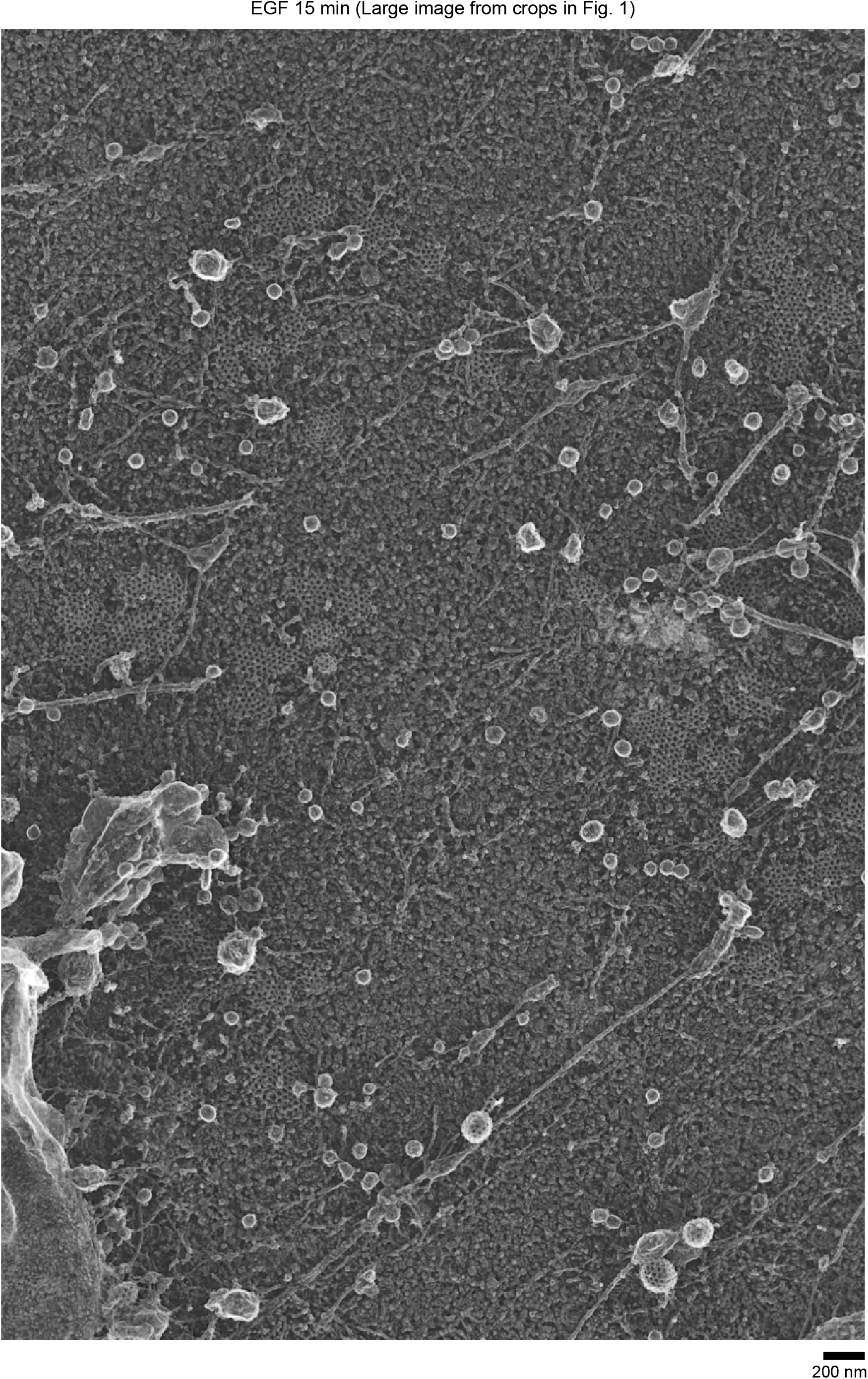
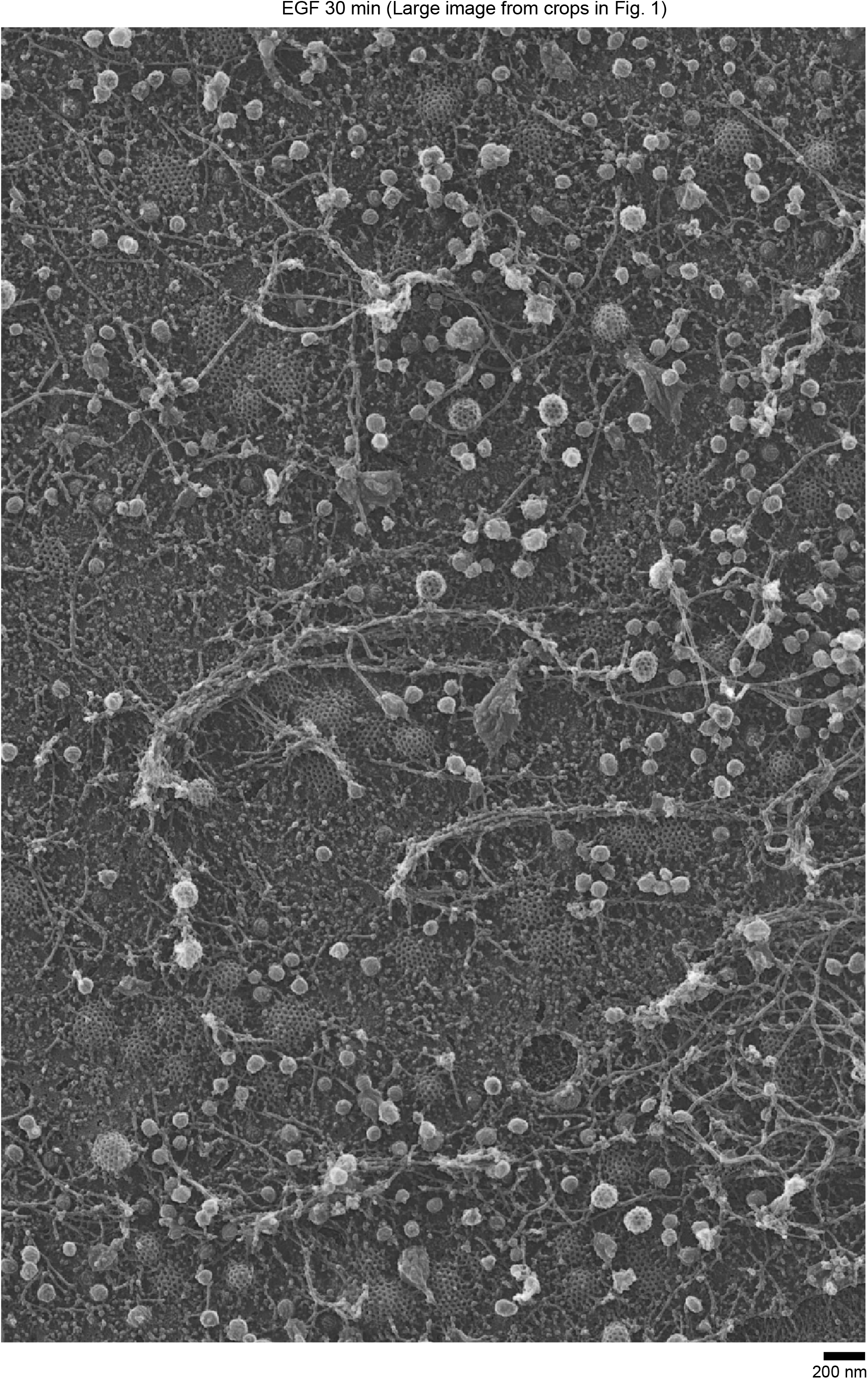
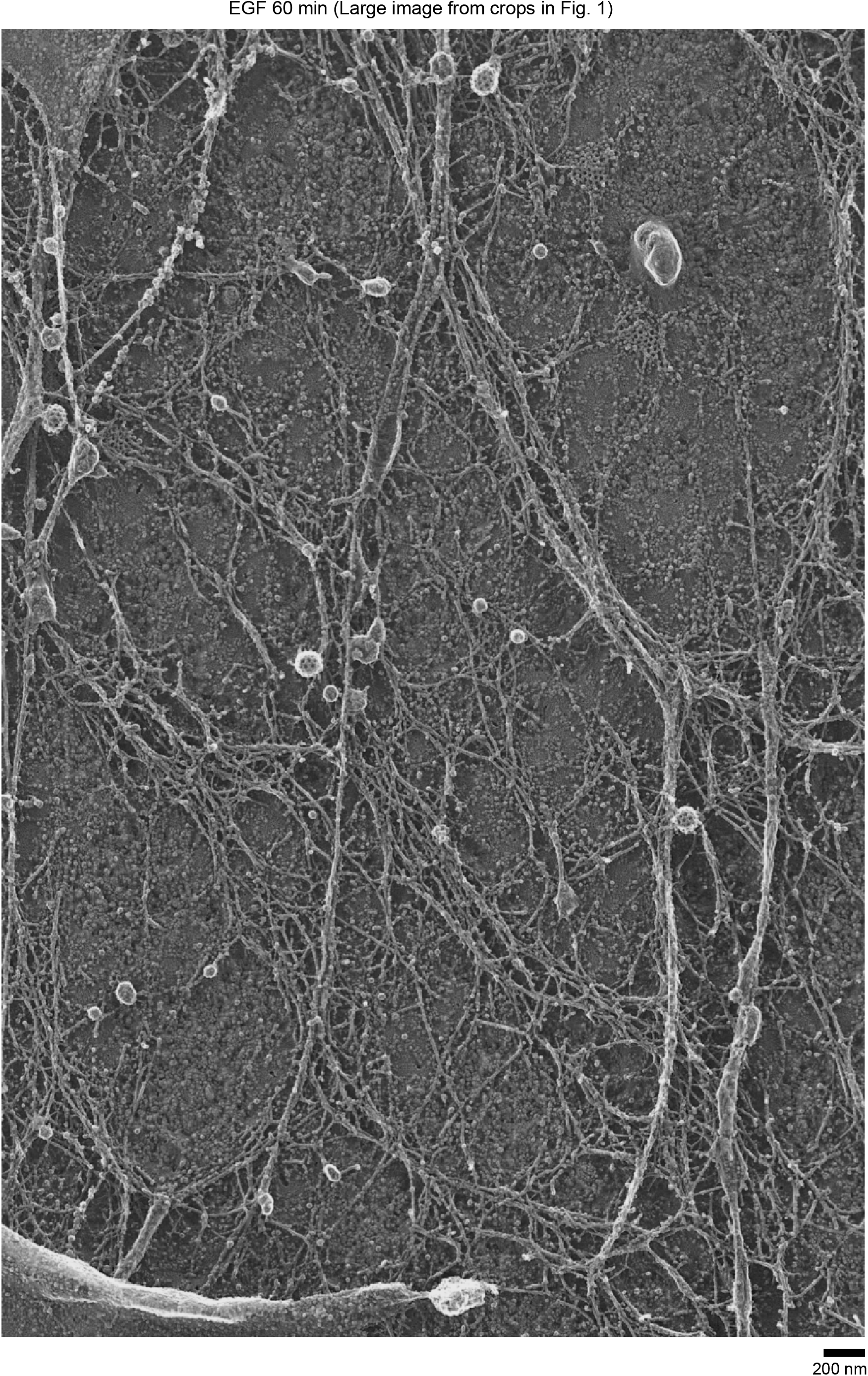
Original PREM images of cells from which the cropped images in Figure 1 were derived. PREM images of control (Ctrl) HSC3-EGFR-GFP cells, stimulated with 50 ng/mL EGF for 0, 2, 5, 15, 30 and 60 min. Scale bars are 200 nm.

**Supplementary Figure 3.**
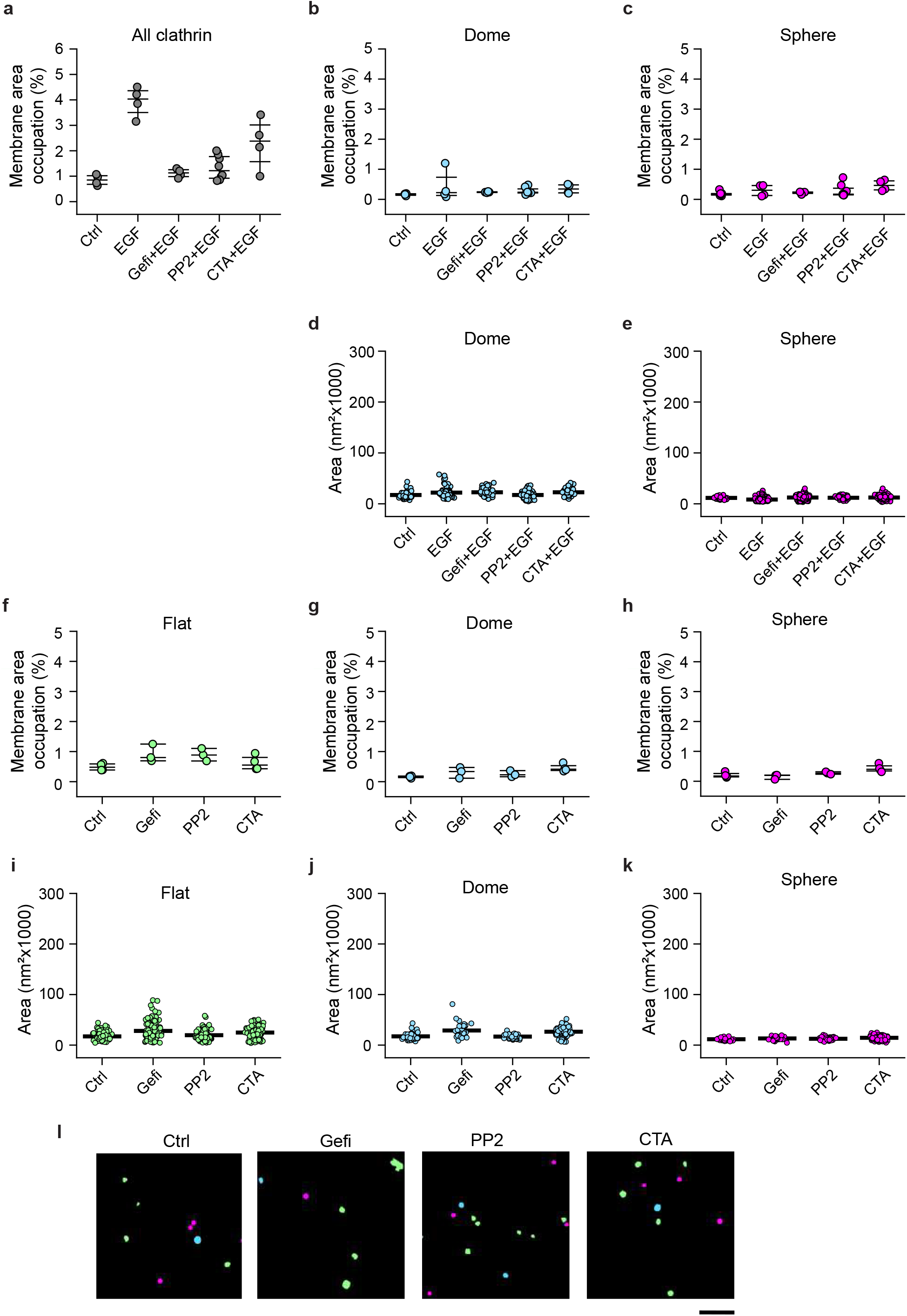
Morphometric analysis of PREM images of cells treated with different drugs. **a-c**, Morphometric analysis of the percentage of plasma membrane area occupation for (**a**) all CCSs, (**b**) dome, and (**c**) sphere structures in control (Ctrl) HSC3-EGFR-GFP cells or treated with 50 ng/mL EGF for 15 min in the absence (EGF) or presence 10 μM gefitinib (Gefi+EGF), 10 μM PP2 (PP2+EGF) and 10 μM cilengitide acid (CTA+EGF). I-shaped box plots show median extended from 25th to 75th percentiles, and minimum and maximum data point whiskers with a coefficient value of 1.5. **d-e**, Morphometric analysis of the size of (**d**) dome and (**e**) sphere clathrin structures in cells treated as in (**a-c**). Dot plots show every structure segmented, the bar indicate the median. **f-h**, Morphometric analysis of the percentage of plasma membrane (PM) area occupation for (**f**) flat, (**g**) dome, and (**h**) sphere structures in control (Ctrl) cells or treated only with the drugs in (**a-c**). **i-k**, Morphometric analysis of the size of (**i**) flat, (**j**) dome and (**k**) sphere clathrin structures in cells treated as in (**f-h**).Ctrl: N_flat_=141, N_dome_=46, N_sphere_=68; N_cells_=4; EGF: N_flat_=559, N_dome_=67, N_sphere_=207, N_cells_=4; Gefi: N_flat_=153, N_dome_=65, N_sphere_=72, N_cells_=4; Gefi+EGF: N_flat_=160, N_dome_=69, N_sphere_=103, N_cells_=4; PP2: N_flat_=109, N_dome_=36, N_sphere_=53, N_cells_=3; PP2+EGF: N_flat_=267, N_dome_=88, N_sphere_=61, N_cells_=4; CTA: N_flat_=171, N_dome_=137, N_sphere_=229, N_cells_=4; CTA+EGF: N_flat_=244, N_dome_=68, N_sphere_=167, N_cells_=4. **i**, Representative masks of segmented cells treated as in (**f-h**). **l**, Representative masks of segmented cells treated as in (**a,f**). Scale bar is Ctrl and EGF data are from Figure 1 and shown for reference.

**Supplementary Figure 4.**
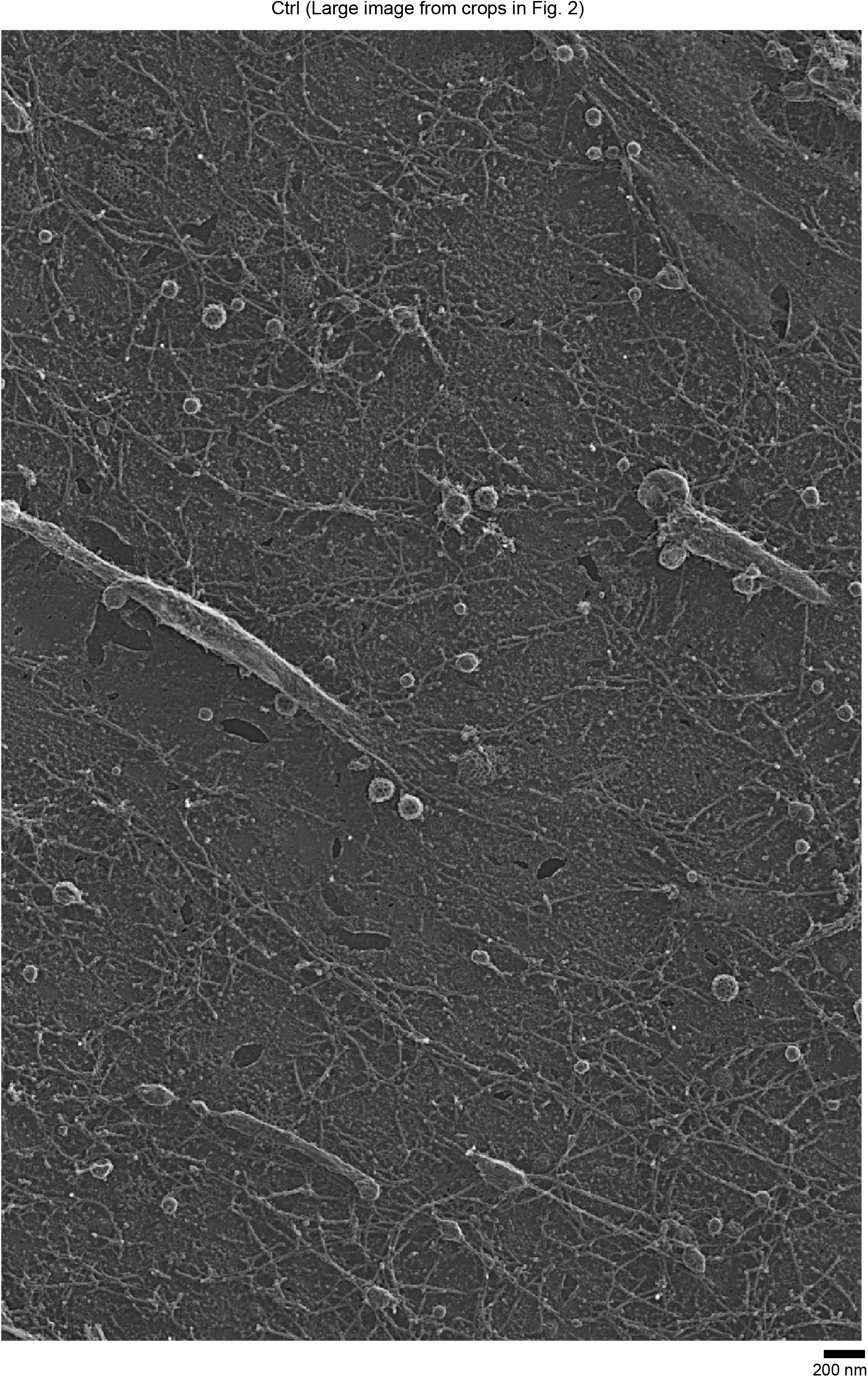
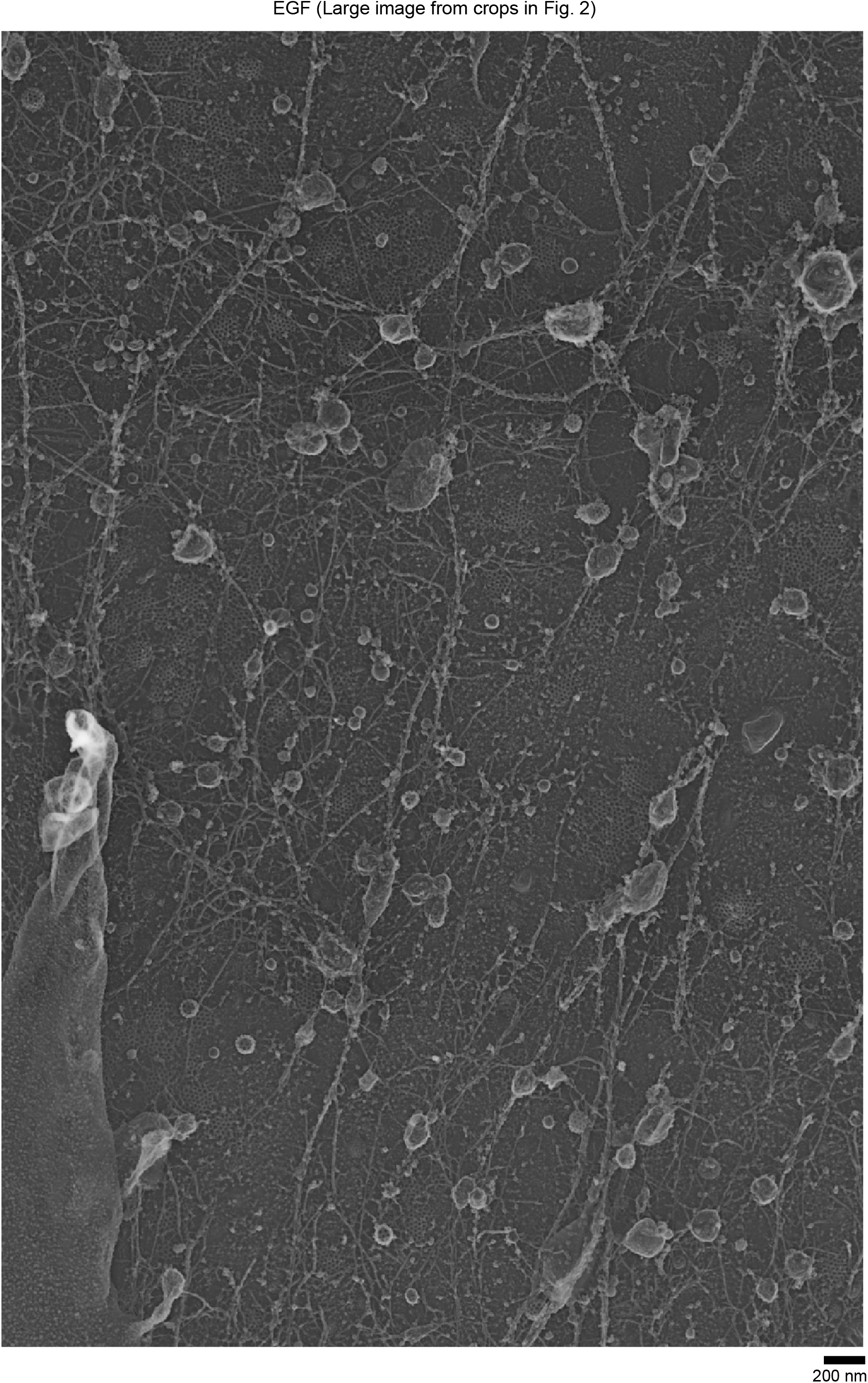
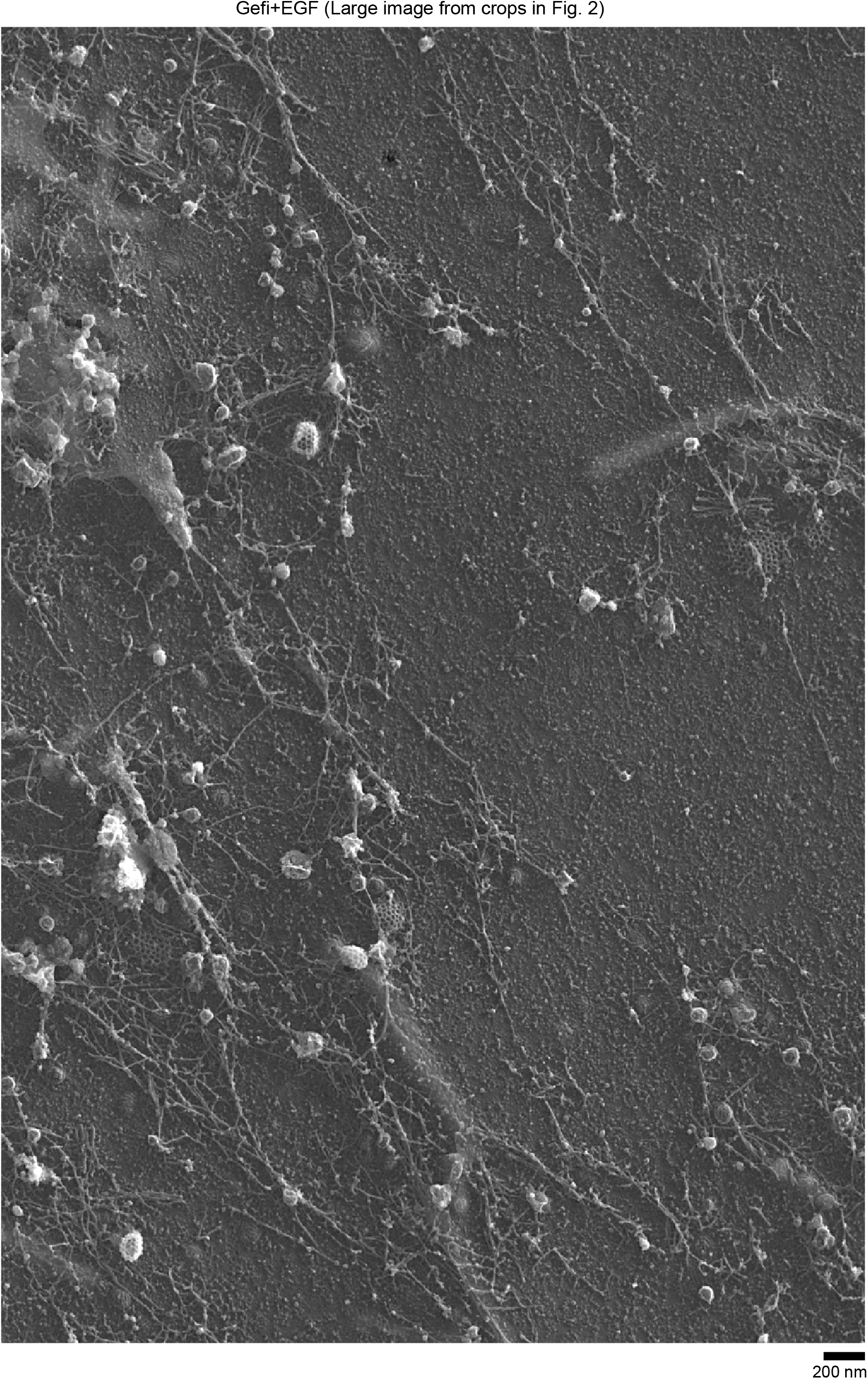
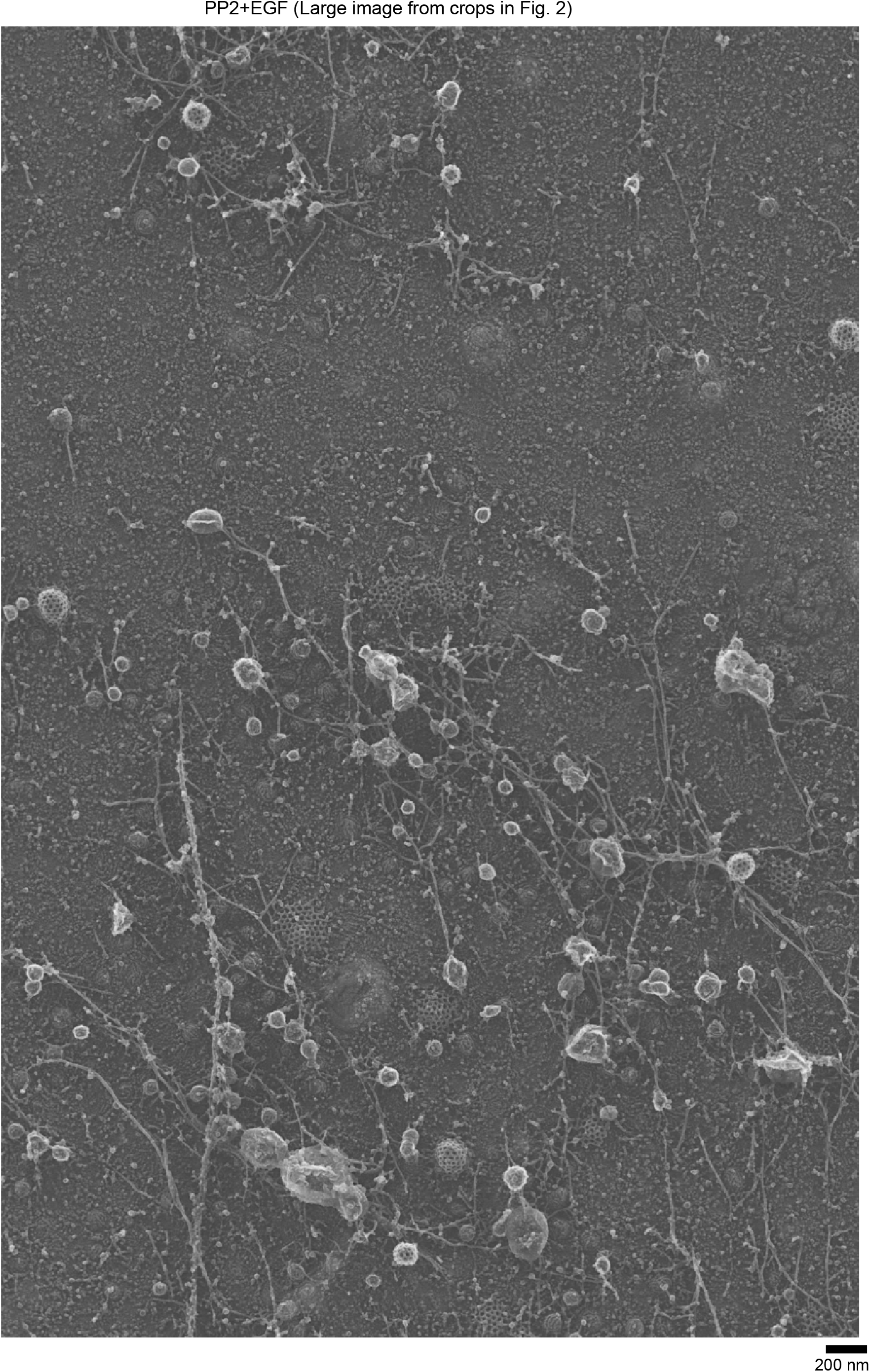
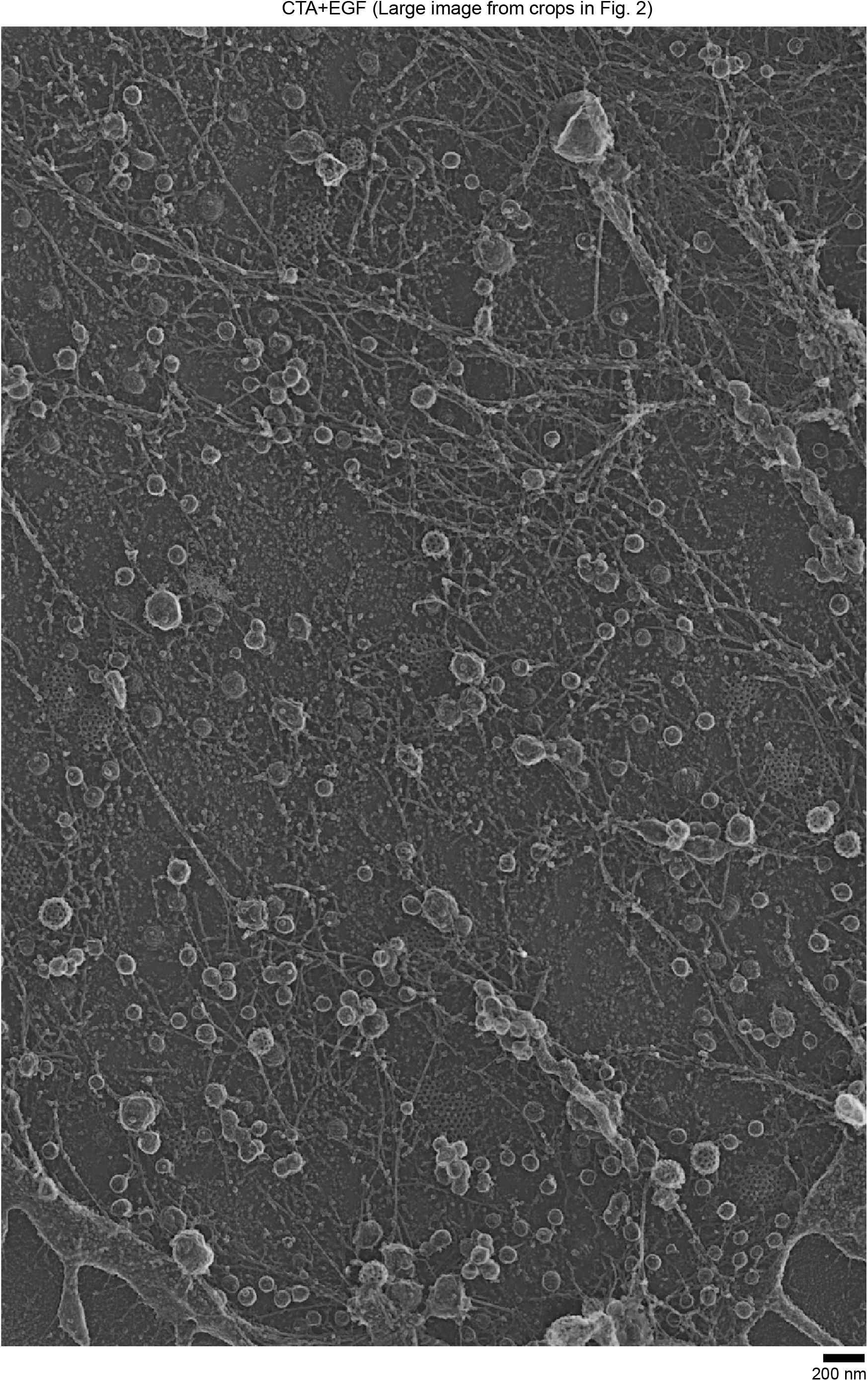
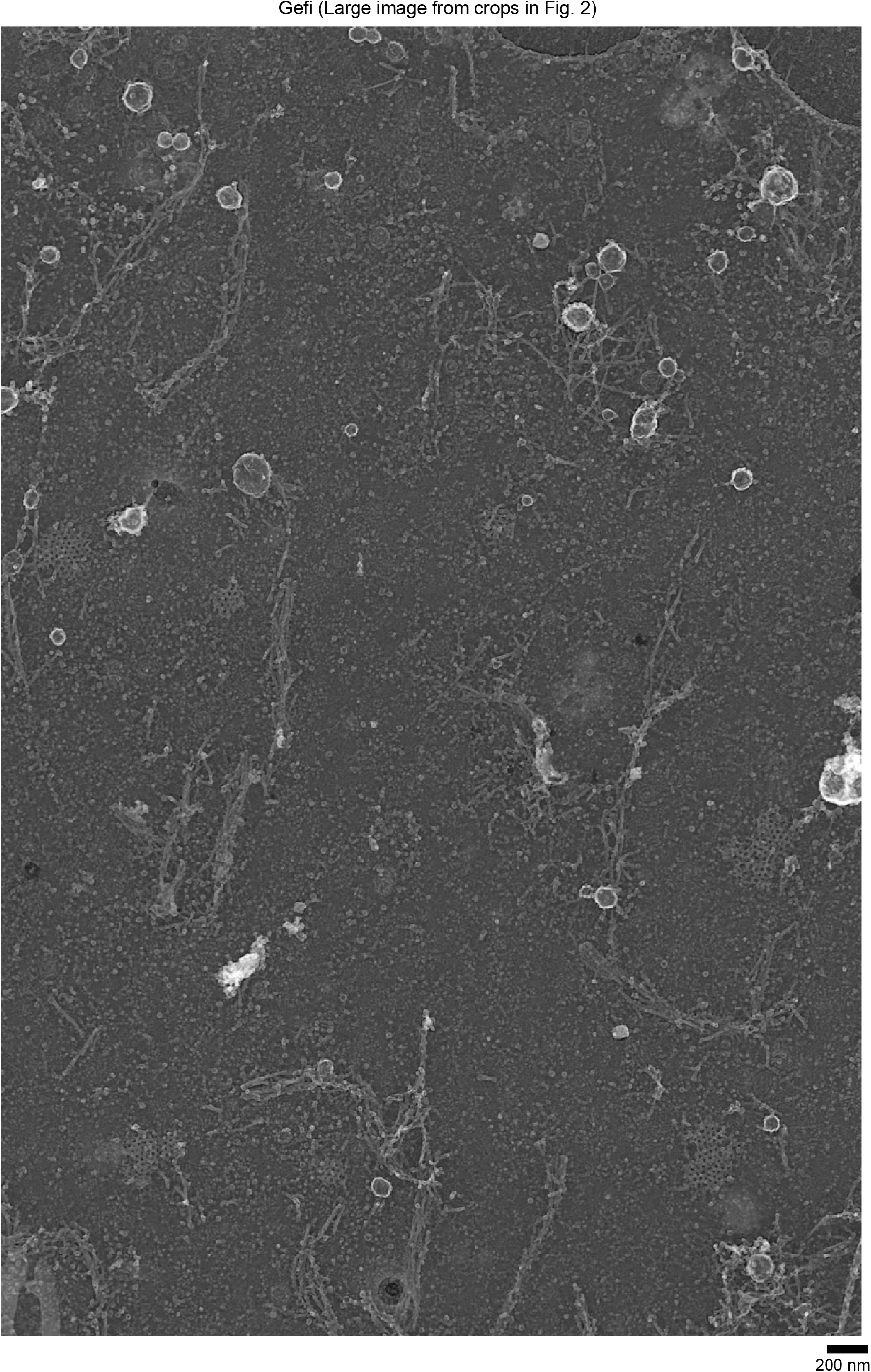
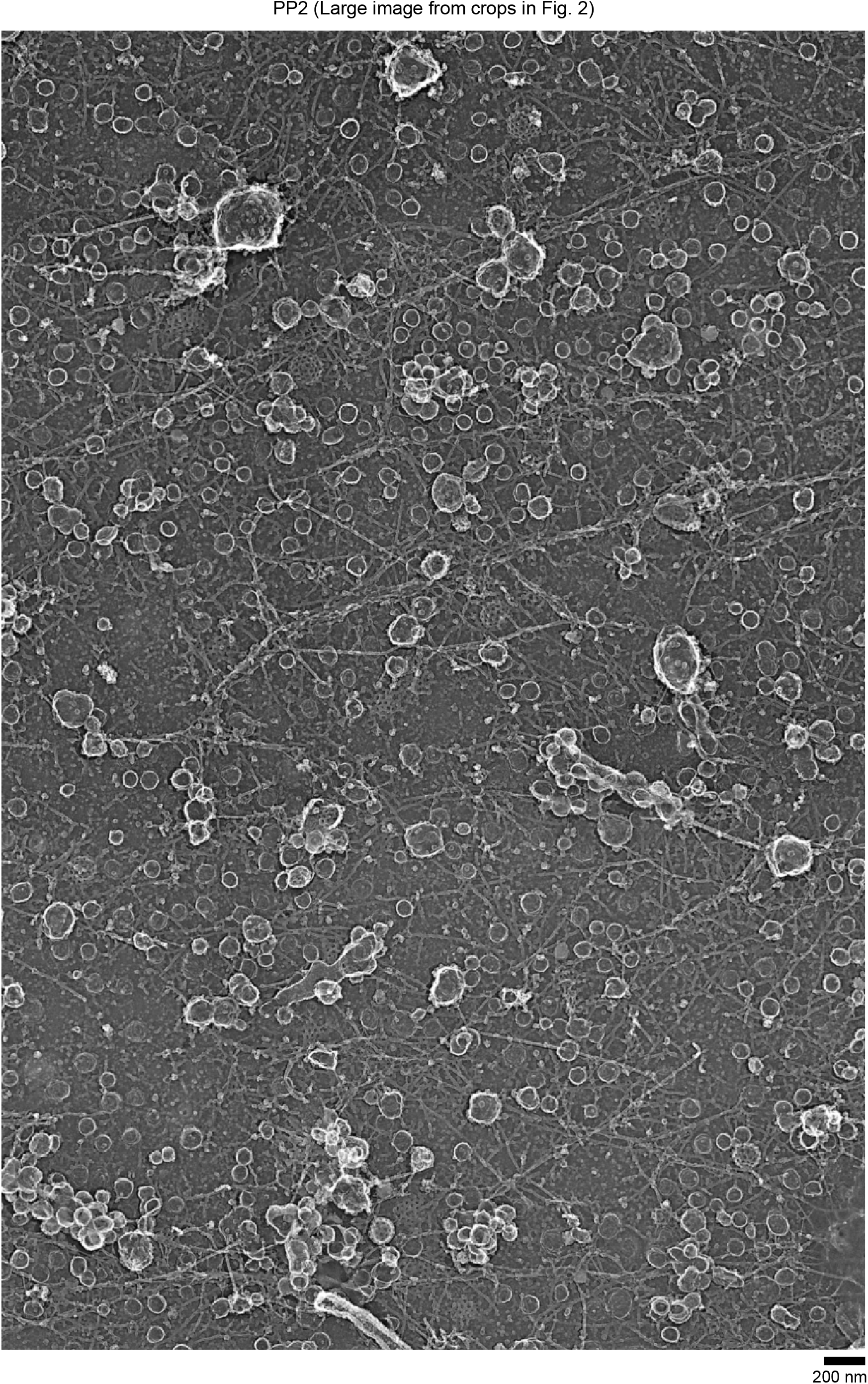
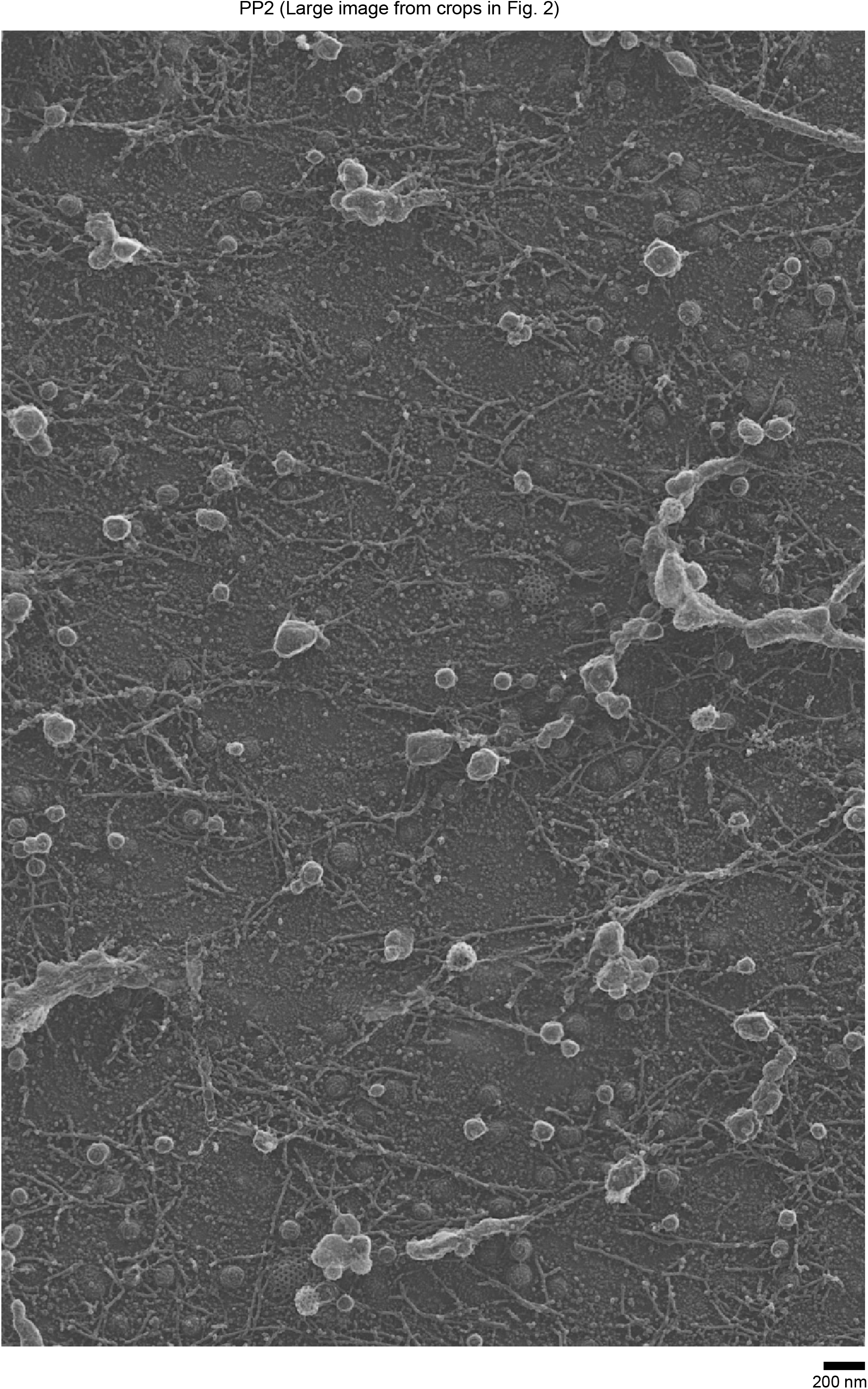
Original PREM images of cells treated with different drugs. Original PREM images of cells from which the cropped images in Figure 2 and masks in Supplementary Figure 2 were derived. Shown are control (Ctrl) HSC3-EGFR-GFP cells or treated with 50 ng/mL EGF for 15 min in the absence (EGF) or presence 10 μM gefitinib (Gefi+EGF), 10 μM PP2 (PP2+EGF), 10 μM cilengitide acid (CTA+EGF) and the drugs alone (Gefi, PP2, CTA). Scale bars are 200 nm.

**Supplementary Figure 5.**
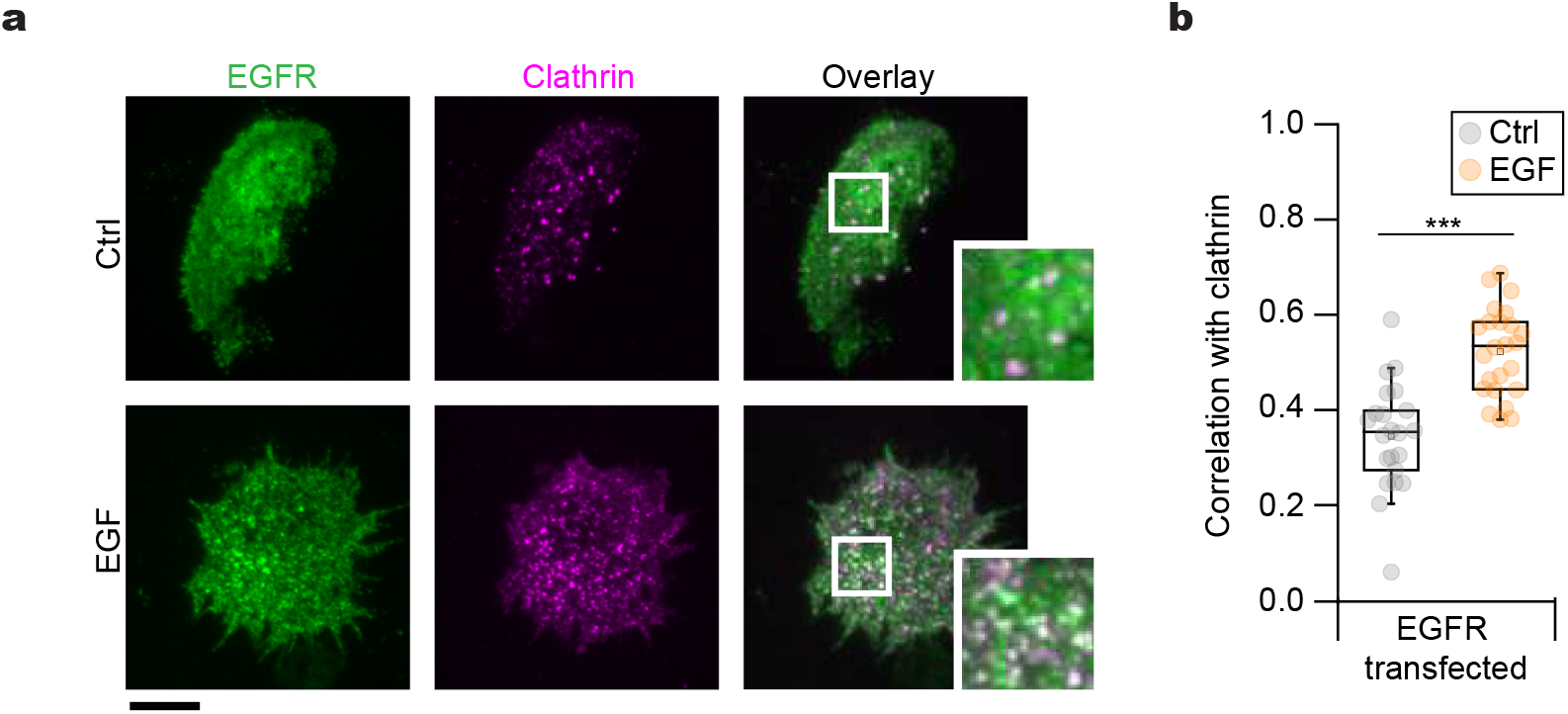
EGFR-GFP correlates with clathrin after EGF stimulation. **a**, Representative TIRF images of HSC3 WT cells co-transfected with mScarlet-CLCa and EGFR-GFP before (Ctrl) or after 50 ng/mL EGF stimulation for 15 min. **b**, Automated correlation analysis of (**a**). Significance was tested by a two-tailed unpaired t-test, \**P*_*EGFR*_=4.2×10^−7^. N_EGFR-Ctrl_= 22 cells – 1148 spots, N_EGFR-EGF_=24 cells – 1305 spots. Dot box plots show median extended from 25th to 75th percentiles, mean (square) and minimum and maximum data point whiskers with a coefficient value of 1.5. Scale bar is 10 μm; insets are 7.3×7.3 μm.

**Supplementary Figure 6.**
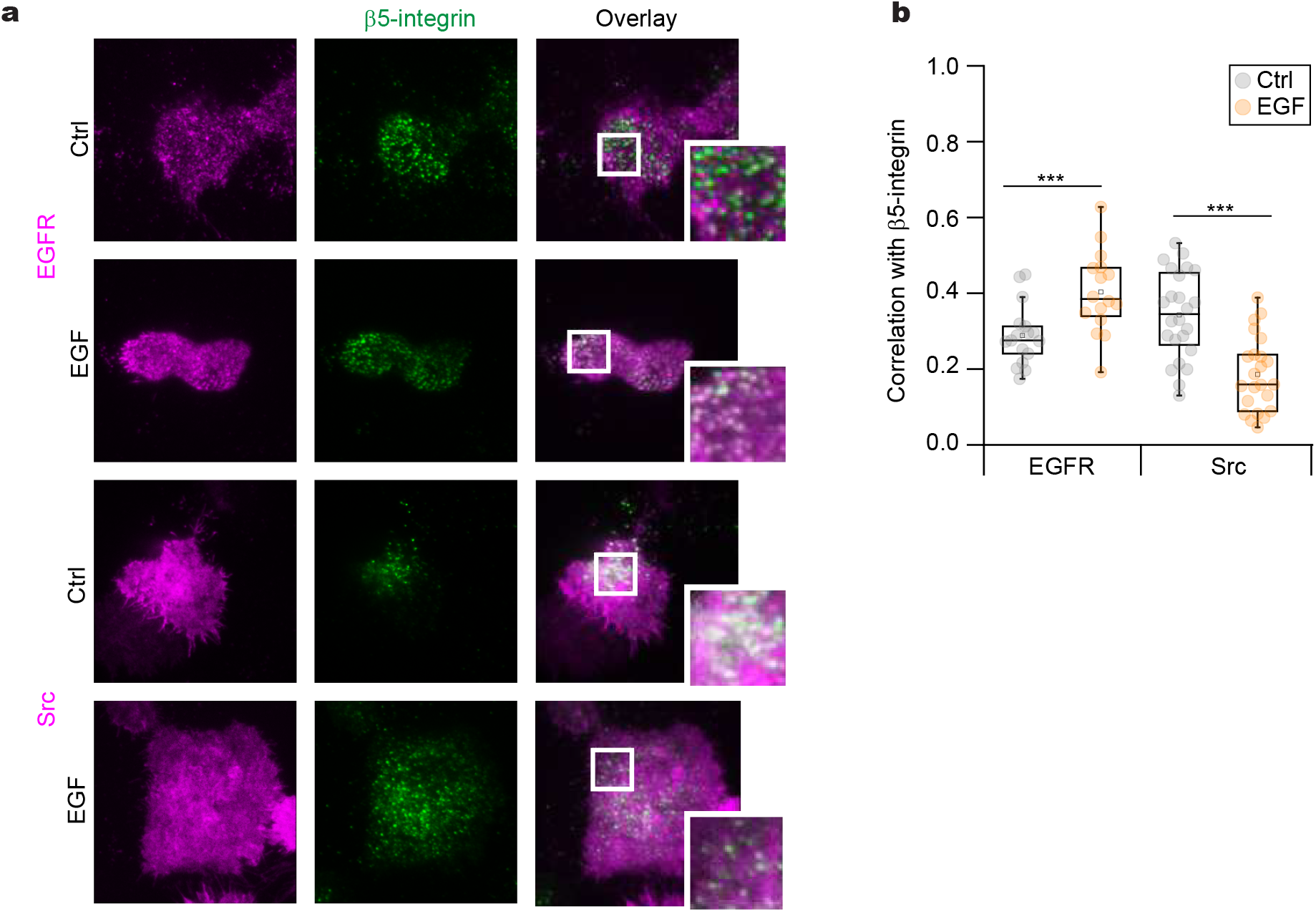
Differential location of EGFR and Src in β5-integrin enriched structures. **a**, Representative TIRF images of HSC3 WT cells co-transfected with β5-integrin-GFP and either EGFR-mScarlet or mCherry-Src before (Ctrl) or after 50 ng/mL EGF stimulation for 15 min. **b**, Automated correlation analysis of (**a**). Significance was tested by a two-tailed unpaired t-test \**P*_*EGFR*_= 8.4×10^−4^, \**P*_*Src*_= 1.2×10^−6^. N_EGFR-Ctrl_=17 cells – 1516 spots, N_EGFR-EGF_=16 cells – 1416 spots, N_Src-Ctrl_= 24 cells – 1936 spots, N_Src-EGF_=23 cells – 1872 spots. Dot box plots show median extended from 25th to 75th percentiles, mean (square) and minimum and maximum data point whiskers with a coefficient value of 1.5. Scale bar is 10 μm; insets are 7.3×7.3 μm.

**Supplementary Figure 7.**
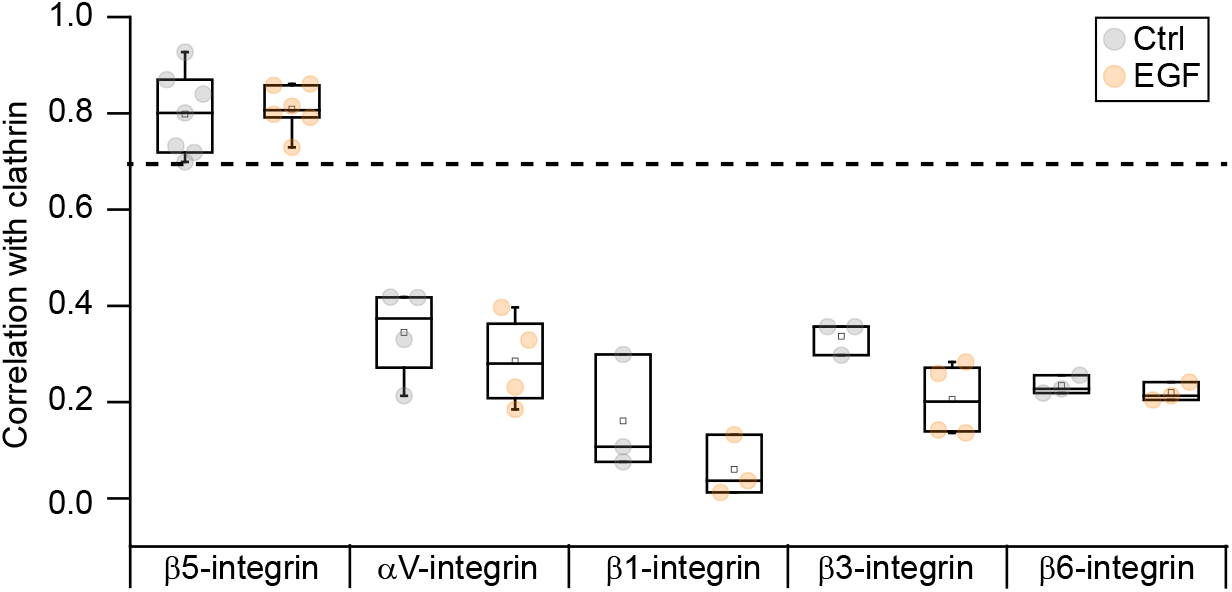
Clathrin coated structures are mainly enriched with β5-integrin. Automated correlation analysis of HSC3 WT cells co-transfected with mScarlet-CLCa and the indicated integrin tagged with GFP before (Ctrl) or after 50 ng/mL EGF stimulation for 15 min. Cell were imaged using TIRFM. Dot box plots show median extended from 25th to 75th percentiles, mean (square) and minimum and maximum data point whiskers with a coefficient value of 1.5. Dots represent the mean correlation value of independent experiments.

**Supplementary Figure 8.**
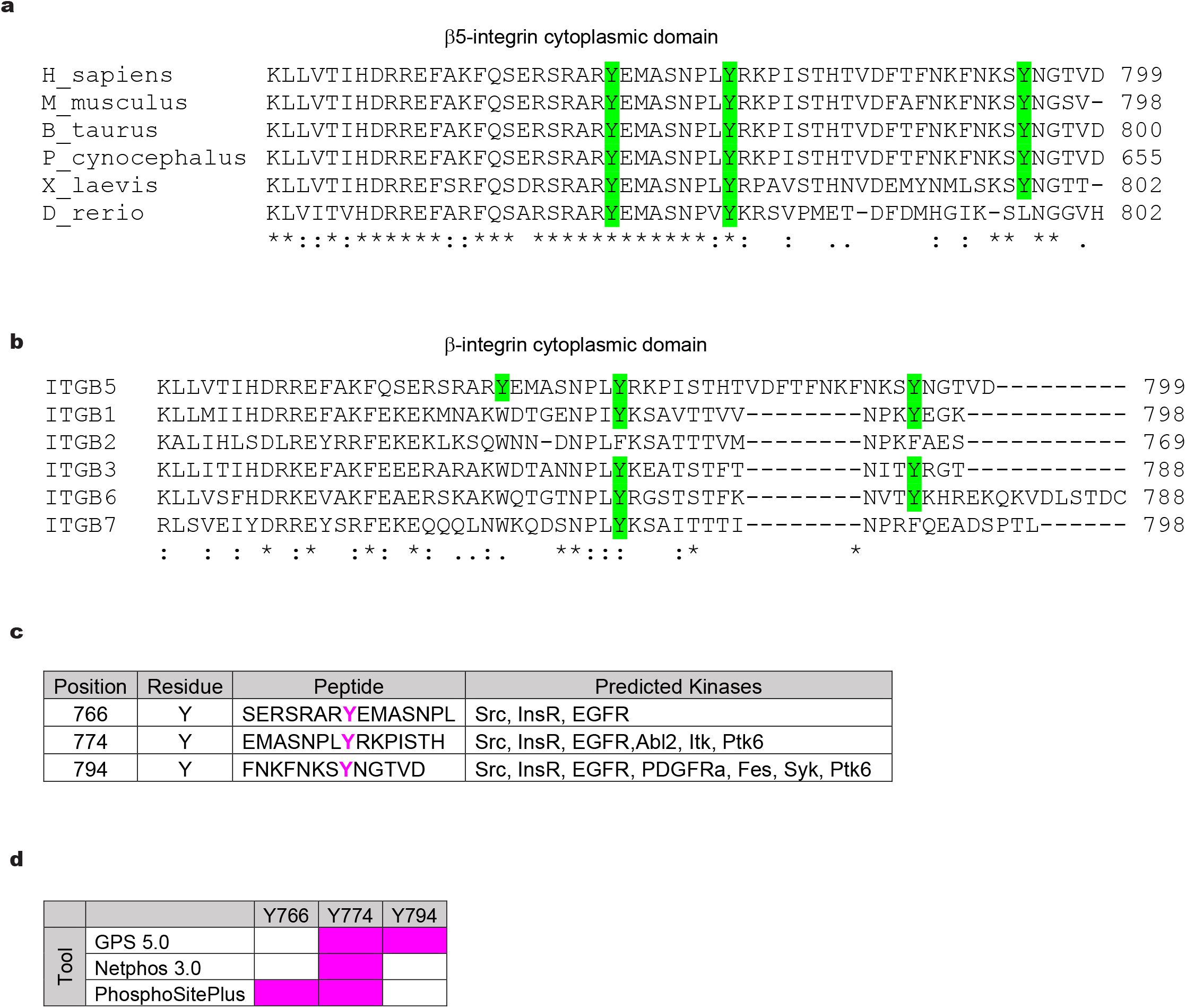
*In silico* analysis of β5-integrin. **a**, Sequence alignment of cytoplasmic domain of different β5-integrin orthologues. Tyrosine residues are marked in green. Symbols: i) **\***, single fully conserved residue; ii):, conservative; iii)., noneconservative. **b**, Sequence alignment of cytoplasmic domain of different β-integrin subfamily members. Symbols as in (a). **c**, Bioinformatic prediction of the possible protein kinases involved in the posttranslational modification of the β5-integrin cytoplasmic domain. The phosphopeptides identified by Netphos 3 and GPS 5 are indicated with the putative modified residues in magenta; the residue position is indicated, as well as the protein kinases most likely involved in the catalysis of the ATP phosho-transfer reaction. **d**, β5-integrin phosphorylation sites prediction. Tyrosine residues present in the β5-integrin cytoplasmic domain are listed with their sequence position indicated. Residues colored with magenta were predicted to be phosphorylated by the indicated bioinformatic tool.

